# Minimally Sufficient Experimental Design using Identifiability Analysis

**DOI:** 10.1101/2023.10.14.562348

**Authors:** Jana L. Gevertz, Irina Kareva

## Abstract

Mathematical models are increasingly being developed and calibrated in tandem with data collection, empowering scientists to intervene in real time based on quantitative model predictions. Well-designed experiments can help augment the predictive power of a mathematical model but the question of when to collect data to maximize its utility for a model is non-trivial. Here we define data as model-informative if it results in a unique parametrization, assessed through the lens of practical identifiability. The framework we propose identifies an optimal experimental design (how much data to collect and when to collect it) that ensures parameter identifiability (permitting confidence in model predictions), while minimizing experimental time and costs. We demonstrate the power of the method by applying it to a modified version of a classic site-of-action pharmacokinetic/pharmacodynamic model that describes distribution of a drug into the tumor microenvironment (TME), where its efficacy is dependent on the level of target occupancy in the TME. In this context, we identify a minimal set of time points when data needs to be collected that robustly ensures practical identifiability of model parameters. The proposed methodology can be applied broadly to any mathematical model, allowing for the identification of a minimally sufficient experimental design that collects the most informative data.

## INTRODUCTION

Mathematical modeling has become ubiquitous in the natural sciences, particularly in public health and pharmacology, as a tool to both understand existing data and make projections about the future.

Specifically, models have been used to retrospectively analyze experimental data, lending insight into the mechanisms underlying the data and suggesting possible strategies that enhance a desired outcome or limit an undesirable one. More recently, models are being developed and calibrated in tandem with data collection, allowing for model-based predictions to inform future experimental design (Eisenberg & Jain 2017; Hu 2004; Kreutz & Timmer 2009; Rajakaruna & Ganusov 2022; Cassidy 2023; Cárdenas et al. 2022; Luo et al. 2022).

Excitingly, in some arenas, such model-based predictions are empowering scientists to intervene in real time. One example is in the adaptive treatment of metastatic castration-sensitive prostate cancer, where personalized model-informed treatment strategies were adjusted based on a patient’s past and current prostate-specific antigen levels (Zhang et al. 2022). Another example emerged during the COVID-19 pandemic, wherein model-suggested strategies were implemented to mitigate disease spread, which in turn necessitated re-calibrating the model as new data emerged (Cassidy 2023; Buchwald et al. 2021).

### Challenges of Model-Informed Experimental Design

One of the key benefits of a good model is the ability to extrapolate from it. However, depending on how the data are structured and when they are collected, a variety of potential models or parameter values can potentially describe the data used to calibrate it. For instance, in (Kareva & Karev 2018), the authors selected four different tumor growth models and evaluated the goodness of fit of each model to a set of available data. While the authors did show that some models fit the data better than others, for several data sets the differences were marginal. Further, if one were to extrapolate tumor growth projections from the various models that well-describe the data, the growth projections would be quite different. Consequently, model-based decisions can vary significantly based on the model used to extrapolate beyond the data. More recently, Harshe and colleagues (Harshe et al. 2023) noted that even for a simple model, such as the classic logistic model, multiple data points need to be sampled in order to uniquely parametrize the tumor growth curve, with the number of necessary points increasing with the amount of noise in the collected data.

Furthermore, to find parameters that can enable a model to have predictive utility, one sometimes needs a “critical” piece of data that allows all the other model pieces to fit into place. One such example was shown in (Kareva et al. 2023), where the authors used a model that connects drug concentration over time to projected levels of target occupancy (TO) in the tumor microenvironment (TME) and parametrized it using published data for anti-PD-1 checkpoint inhibitor pembrolizumab (Lindauer et al. 2017). The authors then used this model to analyze potential criteria for efficacious dose selection for a different compound targeting a co-expressed target TIGIT. They showed the difference between the doses that achieve full TO in the plasma as compared to doses that achieve full TO in the TME, and used the model to re-discover doses that were in fact taken forward in the clinic. A model that does not reasonably capture the actual relationship between plasma concentration of the drug and its level of target engagement in the TME would not have been able to make this assessment.

As these examples show, not all data are created equal. In order to parametrize a model that can enable decision making, it is critical to have both the correct data type (Kareva et al. 2023) and a sufficient number of data points (Harshe et al. 2023). Unfortunately, not all data can be easily collected due to financial, logistical, or technical reasons, as is the case for invasive procedures, such as biopsies. Therefore, it is particularly important to select appropriate and sufficient data for parametrization if a model is to be used to guide decision making.

### Prior Work on using Identifiability for Model-Informed Experimental Design

Identifiability analysis allows one to rigorously study if model parameters can be uniquely determined from available experimental data. A model is considered structurally identifiable if parameters can be uniquely determined given perfect data (Eisenberg & Jain 2017), a condition which of course can never be satisfied in practice. However, structural identifiability is a necessary, though not sufficient, condition for practical identifiability, which answers this question in the context of real and noisy data.

Practical identifiability analysis has been used to improve experimental design by suggesting measurements that need to be collected to resolve parameter non-identifiability issues given some pre-existing data (Raue et al. 2009). For instance, using a model of tumor spheroid growth under treatment with taxol, practical identifiability analysis revealed that an experiment that measures either the maximum rate of drug-induced death or the drug half-saturation constant is sufficient to resolve parameter non-identifiability issues and consequently to increase confidence in model projections (Eisenberg & Jain 2017). In a mathematical model of ligand binding and trafficking, the authors show how in addition to having measurements of extracellular and bound ligand concentrations, the absolute concentration of one pathway species and intracellular ligand concentrations are needed to make model parameters practically identifiable (Raue et al. 2010).

When this analysis is used to determine the most informative targets and time points for the new measurement, this is referred to as optimal experimental design. Typically, this entails identification of an additional measurement (or set of measurements) that contains maximal information for a parameter of interest (Wieland et al. 2021). For instance, using a model describing lactation in cattle, an assessment of practical identifiability was used to discover four time samplings across 100 days that provide high information content for estimating model parameters (Muñoz-Tamayo et al. 2018). In a mathematical model of a gene regulatory network, the most informative experimental conditions were established using a practical identifiability analysis (Steiert et al. 2012). It is of note that methods that do not use identifiability analysis have also been employed to select time points for measurement in an optimal way (Hu 2004; Kutalik et al. 2004; Cho et al. 2020).

### A Novel Approach to Model-Informed Experimental Design

Herein, we propose a workflow for utilizing practical identifiability analysis as a tool for experimental planning. The model-informed method seeks to determine both the minimal number of experimental measurements needed for a quantity of interest, and when those measurements must be collected, in order to “trust” predictions of a data-calibrated mathematical model. The methodology requires developing a model and using it to create simulated data for a variable of interest such that “complete” simulated data result in the parameters of interest being practically identifiable. We then proceed to find the minimal amount of data needed, and when these data must be collected, to ensure that the parameters remain practically identifiable.

The paper is organized as follows. In the Methods section we review the well-established profile likelihood method for assessing practical identifiability of parameters given available data. We also introduce a modification of the classic pharmacokinetic/pharmacodynamic (PKPD) model with a tumor microenvironment compartment, where drug distribution into the tumor, as well as target synthesis in the TME are not static but are a function of tumor volume. This model, which we calibrate and validate using available experimental data on pembrolizumab, will serve as a case study to demonstrate the execution and applicability of the proposed minimal experimental design methodology. In the Methods section, we also introduce in detail the proposed methodology for model-driven discovery of a minimally sufficient experimental design. In the Results section we use the proposed methodology to recommend a minimal protocol for collecting data on the experimental variable of interest (in this case, percent target occupancy in the TME, which is assumed to drive efficacy). In the Discussion section, we address both the challenges in employing the method, and the benefits of this iterative experimentation-modeling approach for the efficient and robust design of experiments.

## METHODS

### Profile Likelihood for Practical Identifiability

Let *p* be the parameter vector for our differential equation model and let 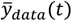 and *σ*^2^(*t*) represent the average and variance, respectively, in the data at time point *t*. For normally distributed measurement noise, the likelihood function is defined as follows (Raue et al. 2013):

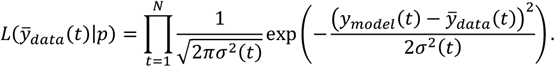

To find the parameter set that gives the best fit to the data, the likelihood function is maximized:

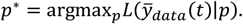

For computational simplicity, the negative of log likelihood function is often minimized instead, which can be shown to be equivalent to minimizing the cost function *ζ* that describes the normalized discrepancy between model predictions and data (Raue et al. 2013):

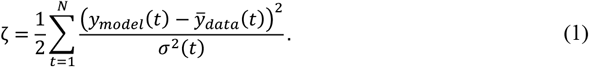

The function *g*(*p*_*i*_) that gives the maximum possible likelihood value for each parameter *p*_*i*_ is called the profile likelihood function. To assess the practical identifiability of model parameters given available data, we will compute the profile likelihood function of each parameter *p*_*i*_ as follows (Raue et al. 2009):

1. Determine a range for the value of *p*_*i*_ from any available theoretical or physiological considerations.
2. Fix 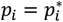, where 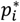 is a value in the range defined in Step 1.
3. For a fixed value of *p*_*i*_ in Step 2, find the parameter set that minimizes the cost function *ζ*.
4. Save the optimal value of the cost function, 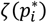.
5. Repeat Steps 2 – 4 for a discrete set of values in the range of parameter *p*_*i*_.
6. Plot *ζ*(*p*_*i*_) to arrive at the profile likelihood function for parameter *p*_*i*_.

The three possible scenarios for the profile of *p*_*i*_ are shown in Figure S1. Figure S1A shows a practically identifiable parameter, as the profile likelihood curve is parabolic with a clear minimum at the optimal value of *p*_*i*_. Further, the range of *p*_*i*_ values within the 95% confidence interval (dashed red line) is finite. Figure S1B represents a parameter that lacks structural identifiability. This is indicated by a flat profile for which an infinite set of parameter values give equally good fits to the data. Figure S1C is indicative of a parameter that is structurally identifiable, but not practically identifiable. While the profile does have a global minimum, it is insensitive to changing the parameter value in one direction.

This can be seen in Figure S1C by observing that the profile does not twice cross its 95% confidence threshold over its domain.

In this work, the profile likelihood method has been implemented in MATLAB® using *ode45* as the numerical differential equation solver with a relative error tolerance of 10^-6^. Parameter fitting is performed using MATLAB’s *fmincon* function with a first-order optimality termination tolerance of 10^-10^. This built-in function executes an interior-point method for solving constrained minimization problems (Nocedal et al. 2014). As we are only fitting a single parameter in all profile likelihood curves generated in this manuscript, the fit parameter is kept on a linear scale. However, we note that when fitting multiple parameters simultaneously, a log scale should be used in the parameter estimation (provided the parameters are non-negative) to avoid potential numerical complications (Raue et al. 2013).

### Case Study: Site-of-Action Model Parametrized for Pembrolizumab

Experiments aimed at characterizing the dose-dependent relationship between drug concentration and tumor size form the backbone of pre-clinical studies in oncology. Typically, the collected time-course measurements are tumor volume and drug concentration in the plasma, which are phenomenologically captured by a simple indirect response model, such as (Simeoni et al. 2004). This correlation might be sufficient for assessing the general dose-response relationship but cannot answer the question of whether the underlying mechanism of action of the drug has been fully engaged.

For this question, we often use pharmacobinding (PB) models (Kareva et al. 2018), which describe the dynamics of the target (such as PD-1 for pembrolizumab), and the reversible binding kinetics between the drug and its target. This allows calculating levels of projected target occupancy, and it is typically expected that if over 90% of the target has been engaged without an effect, then the target may not be a correct one for the selected indication (Kareva et al. 2018). Such PK-PB models can facilitate developing a mechanistic understanding of the dose-response relationship between the drug and the tumor size.

A step further can be taken with site-of-action models (Chudasama et al. 2015; Tiwari et al. 2016; Tiwari et al. 2017) that take into account the drug-target dynamics not only in the plasma but also, as the name suggests, at the site of action, such as the tumor microenvironment. Such models can vary in degrees of complexity from more mechanistic (Betts et al. 2019) to more detailed physiologically based pharmacokinetic models (Zhao et al. 2011; Sager et al. 2015). While such models can be used to calculate projected levels of target occupancy in the TME, it is unclear whether these estimates are truly reliable without actually sampling the TME, the question we will be addressing here.

For that, we developed a modified version of a two-compartment site-of-action model, which describes drug concentration over time in the central (plasma), peripheral and TME compartments. We assume that pharmacobinding occurs in the plasma and TME compartments; while it is possible that some drug-target dynamics occur in the peripheral compartment as well, we assume that it is either negligible with regards to overall dose-response dynamics or cannot be measured; these assumptions can be relaxed if needed.

The model has a standard structure in the plasma compartment, with assumption of intravenous drug administration that is cleared at a rate *k*_10_; the drug distributes to the peripheral compartment at a rate (*V*_1_/*V*_2_)*k*_12_ and back at a rate (*V*_2_/*V*_1_)*k*_21_, where *V*_1_ is the volume of distribution in the central compartment, and *V*_2_ is the volume of distribution in the peripheral compartment. We assume that the free target *T*_*p*_ is synthesized in the plasma at a rate *k*_*syn*_ and, since the model is calibrated to pembrolizumab data whose target PD-1 is membrane-bound, we assume that it is cleared primarily through internalization at a rate *k*_*intP*_. We also assume reversible binding kinetics between the drug and its target, with the drug-target complex in the plasma forming at a rate *k*_*on*_, dissociating at a rate *k*_*off*_, and clearing at the rate *k*_*intP*_.

The PK-PB dynamics in the TME compartment are largely similar with several proposed modifications. Firstly, we assume that the rate of drug distribution into the TME is not constant but a function of the tumor volume, namely, (*V*_1_/(*x* + δ))*k*_1*T*_, where *x* is tumor volume and δ is introduced to prevent division by zero in the limiting case, where the tumor volume tends to zero. We propose that while *V*_1_ and *V*_2_ are treated as constant volumes of distribution (as is standard), the volume of distribution into the tumor be treated as variable, thereby capturing higher or lower distribution of the drug into the TME depending on tumor size. As a consequence of this assumption, we further propose that the rate of target synthesis in the tumor is not constant or at equilibrium as would likely be in the plasma or non-disease compartment, but instead is treated as a function of tumor size. In particular, we assume this rate increases according to a saturating function 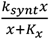, where *k*_*synt*_ is the rate of target synthesis in the tumor (which is likely higher than in the plasma), and *K*_*x*_ is the half-maximal concentration of free target *T*_*TME*_ in the tumor microenvironment.

Additionally, we hypothesize that the apparent rate of drug-target binding in the TME is not necessarily the same as in the plasma, i.e., that *k*_*onT*_ may be different from *k*_*on*_. That said, we expect that once the drug-target complex has been formed, the dissociation rate *k*_*off*_ will remain the same, as that is more likely to be an intrinsic property (Dunlap & Cao 2022). Finally, we assume that the tumor grows logistically and is killed as a function of the percent target occupancy in the tumor, which is calculated as 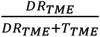, where *DR* _*TME*_ is the concentration of the drug-target complex in the tumor and *T*_*TME*_ is the free target in the TME.

The resulting system of equations is as follows:

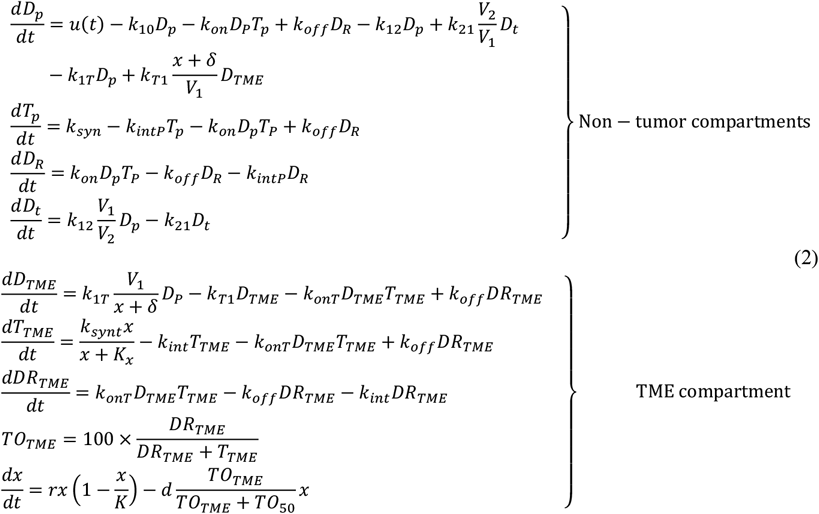

The structure of the model is summarized in Figure 1. Variable definitions, initial conditions, and calibrated parameter values are summarized in Table 1.

**Table 1.** Parameter values used in System (2).

| Variable | Description | Initial Condition | Reference |
| --- | --- | --- | --- |
| $D_p$ | Concentration of drug in plasma (mg/L) | $D_p(0) = 0$ | n/a |
| $T_p$ | Concentration of unbound (free) drug target in plasma (nM) | $T_p(0) = 10$ | (Lindauer et al. 2017) |
| $D_R$ | Concentration of drug-target complex in plasma (nM) | $D_R(0) = 0$ | n/a |
| $D_t$ | Concentration of drug in peripheral (mg/L) | $D_t(0) = 0$ | n/a |
| $D_{TME}$ | Concentration of drug in TME (mg/L) | $D_{TME}(0) = 0$ | n/a |
| $T_{TME}$ | Concentration of unbound (free) drug target in TME (nM) | $T_{TME}(0) = 43$ | (Lindauer et al. 2017) |
| $DR_{TME}$ | Concentration of drug-target complex in TME (nM) | $DR_{TME}(0) = 0$ | n/a |
| $x$ | Tumor volume (mm<sup>3</sup>) | $x(0) = 38$ | fit |
| Parameter | Description | Value | Reference |
| $V_1$ | Volume of distribution, plasma compartment (mL/kg) | 70 | Calibrated by fitting model to data digitized from (Lindauer et al. 2017) |
| $V_2$ | Volume of distribution, peripheral compartment (mL/kg) | 33 | |
| $k_{10}$ | Rate of clearance from plasma compartment (d<sup>-1</sup>) | 5/70 | |
| $k_{on}$ | Binding rate of drug to target in plasma (nM<sup>-1</sup>d<sup>-1</sup>) | 0.005 | |
| $k_{off}$ | Dissociation rate of drug-target complex (d<sup>-1</sup>) | 1.35e-4 | |
| $k_{12}$ | Rate constant for drug distribution from plasma to peripheral compartment (d<sup>-1</sup>) | 22/70 | |
| $k_{21}$ | Rate constant for drug distribution from peripheral to plasma compartment (d<sup>-1</sup>) | 22/33 | |
| $k_{1T}$ | Rate constant for drug distribution from plasma to tumor microenvironment (d<sup>-1</sup>) | 0.3 | fit |
| $k_{T1}$ | Rate constant for drug distribution from tumor microenvironment to plasma compartment (d<sup>-1</sup>) | 0.3 | fit |
| $\delta$ | Numerical correction term to avoid division by zero (mm<sup>3</sup>) | 1e-4 | n/a |
| $k_{intP}$ | Decay or internalization rate of drug target in plasma (d<sup>-1</sup>) | 4.4 | fit |
| $k_{syn}$ | Rate of target synthesis in plasma (nM/d) | 44 | Computed: = $T_p(0)k_{intP}$ |
| $k_{onT}$ | Binding rate of drug to target in TME (nM<sup>-1</sup>d<sup>-1</sup>) | 0.01 | fit |
| $k_{synt}$ | Maximum rate of target synthesis in plasma (nM/h) | 14190 | fit |
| $k_{int}$ | Decay or internalization rate of drug target in TME (d<sup>-1</sup>) | 4.4 | fit |
| $r$ | Intrinsic tumor growth rate (d<sup>-1</sup>) | 0.148542 | fit |
| $K$ | Tumor carrying capacity (mm<sup>3</sup>) | 10000 | n/a |
| $d$ | Maximum rate of tumor kill by pembrolizumab (d<sup>-1</sup>) | 0.38 | fit |
| $TO_{50}$ | Percent target occupancy in TME that results in 50% kill rate by pembrolizumab | 43 | fit |

**Figure 1.**
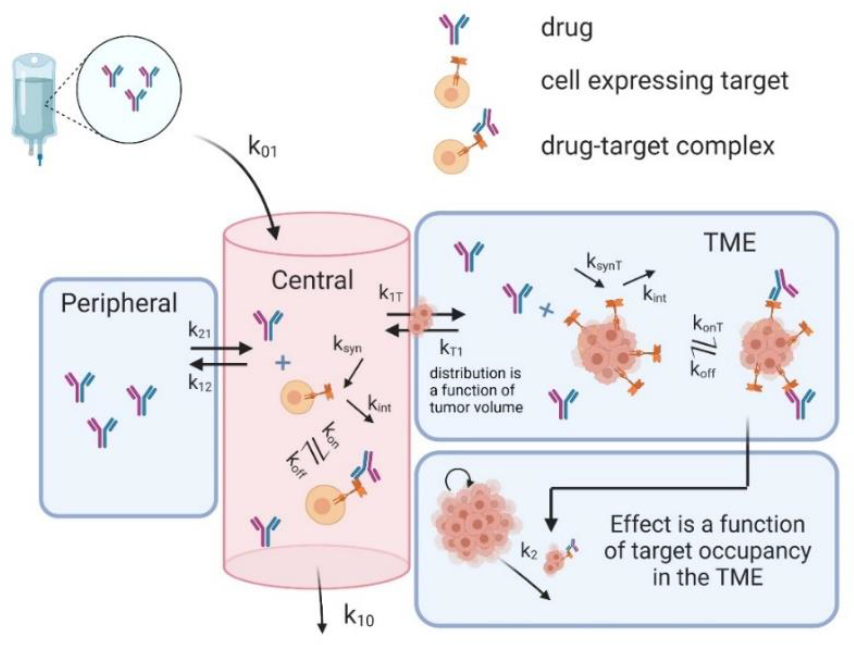
The diagram of model described by System (2).

The model was calibrated to digitized PK data (Figure 2A) for pembrolizumab reported in (Lindauer et al. 2017) and tumor growth inhibition (TGI) data reported in (Agency 2015). The reason this particular PK dataset was chosen is that it includes measurements of percent TO in the TME, which is typically not available. TGI curves in (Agency 2015) are measured for 2 mg/kg and 10 mg/kg of pembrolizumab, administered on average 3.5 days apart, for C57BL/6 mice implanted with MC38 syngeneic colon adenocarcinoma cells. We further calibrated model parameters to fit the PK-TGI relationship for the dose of 10 mg/kg (Figure 2B). We also report the projected levels of percent TO in plasma as compared to the TME (Figure 2C) to emphasize the importance of capturing drug-target dynamics in the TME, as this is where it is expected to drive efficacy.

**Figure 2.**
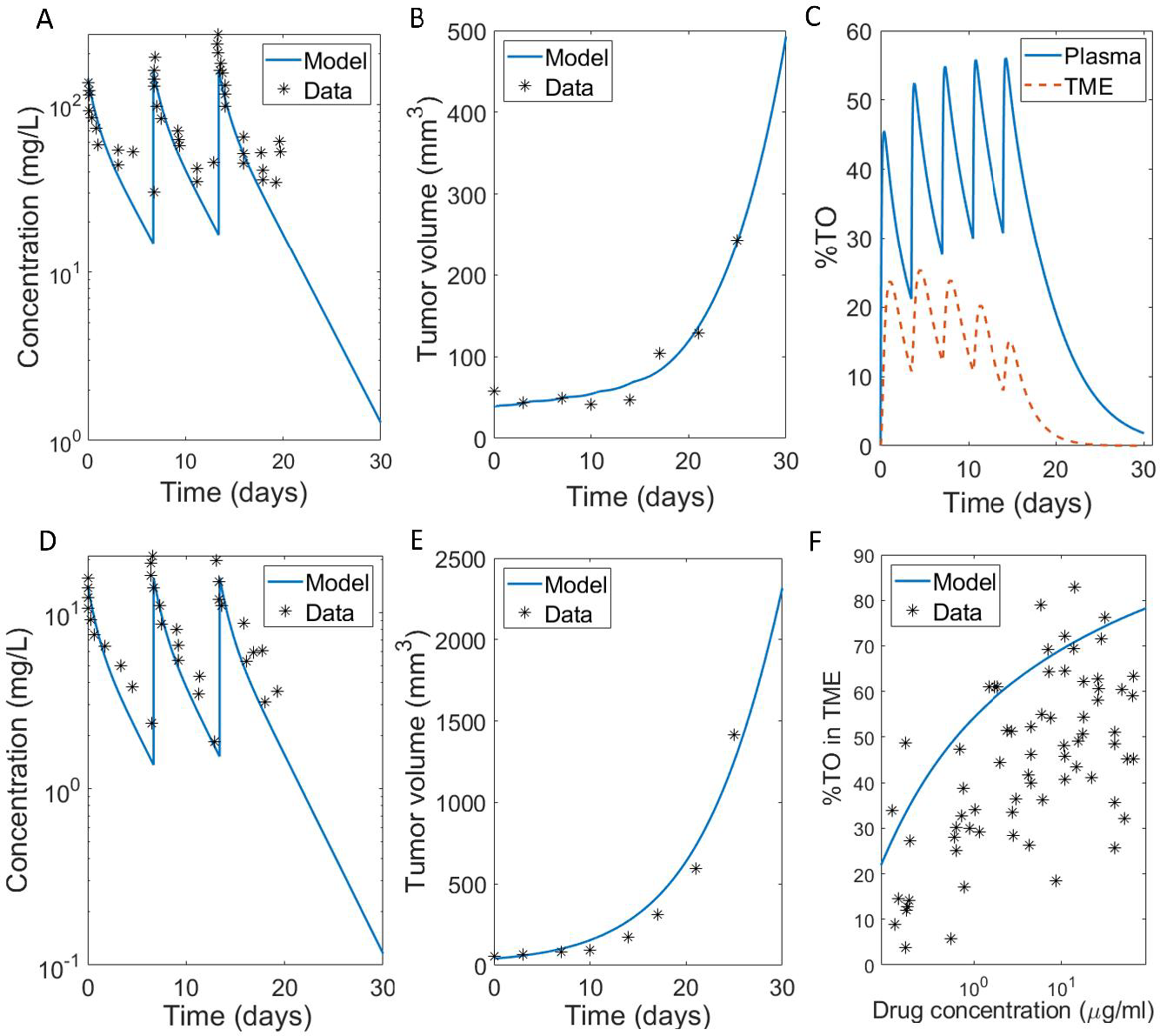
(A)-(B) Calibrated fits of model in System (2) to (A) PK curves for three doses of 10 mg/kg of pembrolizumab given weekly (Lindauer et al. 2017), (B) TGI data for five doses of 10 mg/kg of pembrolizumab given on average every 3.5 days (Agency 2015). (C) Corresponding projected levels of %TO in plasma and the TME. (D)-(E) Validation of model in System (2) on untrained data. (D) PK curves for three doses of 1 mg/kg of pembrolizumab given weekly (Lindauer et al. 2017), (E) TGI data for five doses of 2 mg/kg of pembrolizumab given on average every 3.5 days (Agency 2015). (F) Percent target occupancy in TME data, with data digitized from (Lindauer et al. 2017).

Model parameterization was validated using untrained data. Figure 2D demonstrates that we were able to successfully recapitulate the PK curves for three doses of 1 mg/kg of pembrolizumab given weekly (Lindauer et al. 2017), and Figure 2E shows that we were able to describe the TGI data for five doses of 2 mg/kg of pembrolizumab given on average every 3.5 days (Agency 2015). We note that, without the %TO in TME data (Figure 2F), there was a large number of parameter sets that could recapitulate the PK and TGI equally well, further emphasizing that this piece of data was critical to model parameterization.

### Designing Minimally Sufficient Experimental Protocol

We propose the following workflow, summarized in Figure 3, for using the profile likelihood method to design a minimally sufficient experimental protocol:

**Figure 3.**
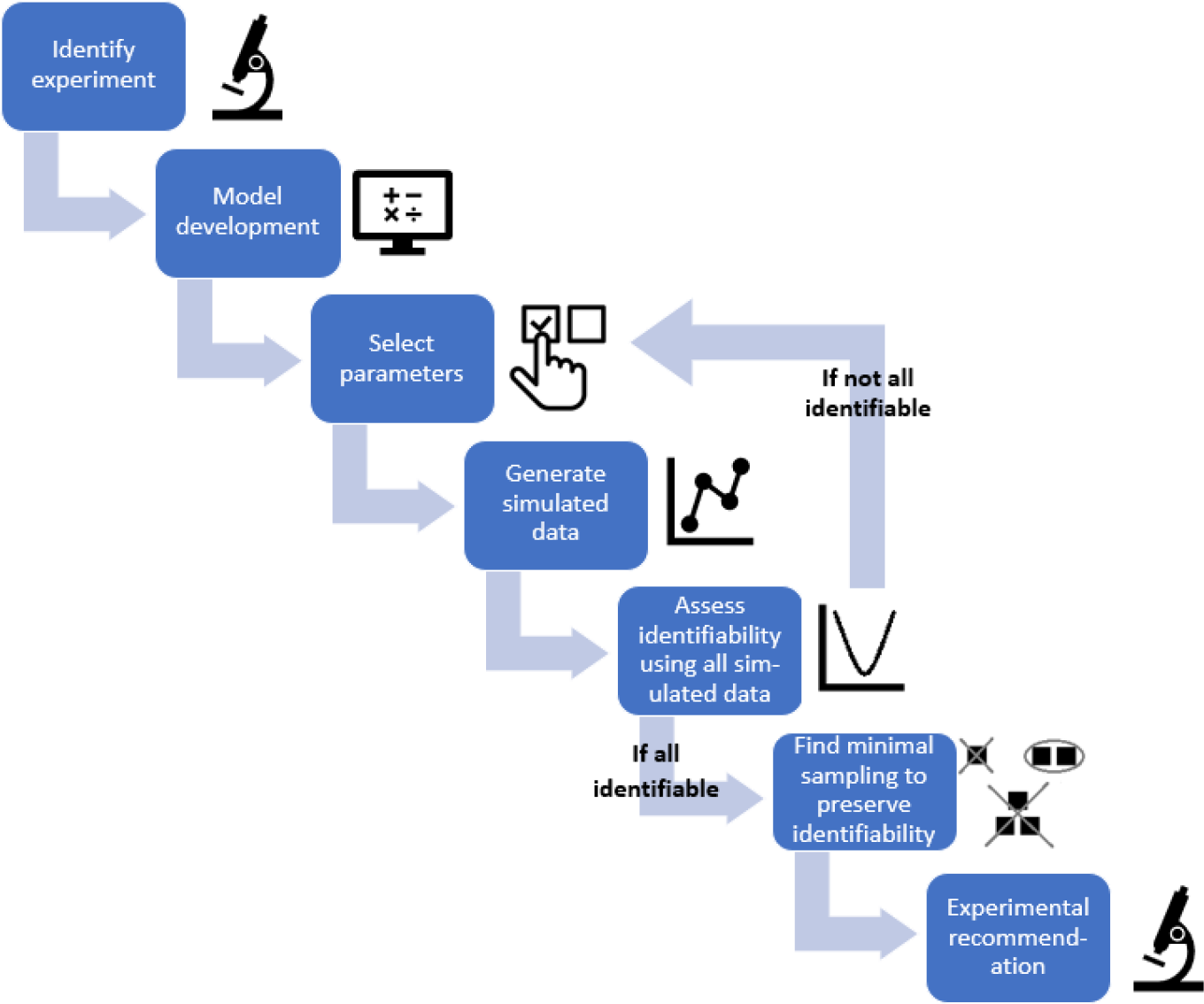
Flowchart of the proposed method of using practical identifiability for model-driven minimal experimental design.

1. *Identify experiment that measures the variable of interest*. We recommend focusing on an experiment that, while potentially expensive and/or invasive, provides data needed to inform decision making from your modeling work.
2. *Model development, parameterization, and validation*. As is always the case in modeling, the validity of its predictions is constrained by the accuracy of the model. Best-practices for designing a fit-for-purpose model have been extensively reviewed elsewhere (Brady & Enderling 2019; Beckman et al. 2020; Sher et al. 2022).
3. *Select parameters of interest*. As fit-for-purpose biological models often contain double digit numbers of parameters, it is often not feasible or desirable to allow all parameters in a model *ℳ* to vary. If *ℳ* has *r* parameters, the goal of this step is to identify a subset of *n* < *r* model parameters for further analysis. This can be done via the following considerations:
  a. Work closely with experimental colleagues, and consult the literature, to identify values for as many parameters as possible.
  b. For those parameters that cannot be easily estimated from experimental data, conduct a sensitivity analysis (Qian & Mahdi 2020; Zhang et al. 2015; Zi 2011). Those parameters the model is least sensitive to should be fixed.
  c. Let ***q*** = (*q*_1_, …, *q*_*m*_) be the set of *m* < *r* model parameters that are fixed based on (a) and (b).
  d. Let ***p*** = (*p*_1_, …, *p*_*n*_) be the set of parameters that have not been fixed in the prior step, where *n* = *r* − *m*. This parameter set will be used in the subsequent analysis.
4. *Generate simulated data for the measurement of interest*. Define an underlying distribution for each parameter in ***p***. Randomly sample *K* values of each parameter from its corresponding distribution and determine the model-predicted response by solving *ℳ*(***q, p***_*k*_, *t*), *k* = 1, …, *K*. From the model-predicted response, extract the value of the variable of interest 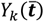 at the time points of interest 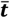. For instance, it is often only feasible to conduct a maximum of one experiment per day, so 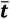 might represent the vector of all days during the experimental time frame; that is, 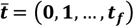.
  a. These distributions could be the output of a fitting procedure, i.e., if nonlinear mixed effects modeling (Owen & Fiedler-Kelly 2014; Olofsen et al. 2004) or Approximate Bayesian Computation (Csilléry et al. 2010; Marjoram 2013) is utilized.
  b. Alternatively, one can assume that each parameter is normally or lognormally (to avoid the possibility of negative parameter values) distributed about its calibrated value. The choice of distribution and its standard deviation should be guided by biological considerations and/or historical data (for instance, how much noise is observed in experimental data), where possible.
5. *Practical identifiability using the data from “complete” simulated experiments*. Use the profile likelihood approach to assess the practical identifiability of all parameters in ***p*** using the simulated data 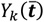 from Step 4. We note that the use of noisy simulated data to explore model identifiability has been considered by others (Wu et al. 2008; Eisenberg & Jain 2017; Raue et al. 2010; Steiert et al. 2012). The goal at this step is for all parameters in ***p*** to be practically identifiable, so if this is not the case, revisit Steps 2 and 3 to right-size your model and reconsider which parameters in the vector ***p*** could be fixed. Proceed to the next step once all parameters in ***p*** are practically identifiable given the simulated data 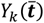.
6. *Determine the minimal number of data points needed to ensure practical identifiability*. Start by asking if it is possible to only conduct *j* = 1 experiment to collect the data on the variable of interest while preserving the practical identifiability of parameters in ***p***. That is, does any 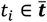 exist for which model parameters ***p*** remain practically identifiable given only the measurement *Y*_*k*_(*t*_*i*_)?
  a. If yes, proceed to Step 7.
  b. If no, repeat the analysis in Step 6 by considering if practical identifiability is ensured if instead you could conduct *j* + 1 experiments to collect data for *Y*_*k*_.
7. *Make minimal experimental design recommendation*. The value of *j* in Step 6 for which practical identifiability of all parameters in ***p*** can be ensured determines the number of experiments that must be conducted and the time when they need to occur. Such a *j* is guaranteed to exist because Step 5 ensured that the parameters in ***p*** are practically identifiable using all the simulated data 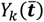. For instance, if *j* = 1 ensured practical identifiability of all parameters in ***p***, only a single experiment needs to be conducted. However, this does not mean that just any single experiment can be conducted. The results in Step 6 can also be used to determine *which subset of experiments* ensure practical identifiability. That is, the method can determine not only the minimal number of experiments to conduct, but when those experiments must be conducted.

We note that while it may appear counterintuitive that the experimental design precedes the modeling step in the proposed workflow, often it is the experimental constraints that will determine the structure of a useful model. For instance, if the only data available to a modeler is TO in the plasma but not in the TME, modeling TO in other compartments will likely not be instructive. In fact, it is likely to introduce a source of additional uncertainty. It is thus the understanding of experimental constraints that guides the development of a practically useful model which can be used to optimize the sampling schedule.

The MATLAB code implementing the methodology for the model described in System (2) is available at https://github.com/jgevertz/minimal_experimental_design. The computational resources and costs of implementing the various stages of the workflow are summarized in Table S1.

## RESULTS

Here we determine how much experimental data on percent target occupancy in the tumor microenvironment is needed to confidently connect PK in the plasma to PD in the TME. That is, what is the minimal experimental protocol that would give us confidence in model predictions? To achieve this goal, we follow the steps set forth above:

1. *Identify experiment that measures the variable of interest*. Here, the variable of interest is percent target occupancy in the TME. This is defined in System (2) as *TO*_*TME*_.
2. *Model development, parameterization, and validation*. We use the validated model in System (2), parameterized using values specified in Table 1.
3. *Select parameters of interest*. We determined that the parameters to be varied in our analyses are the rate of drug-target complex formation in TME, *k*_*onT*_, and the rate of target synthesis in TME, *k*_*synt*_. We arrived at this choice by first removing parameters from consideration that are relatively easy to measure experimentally. This eliminated the PK parameters associated with pembrolizumab, which are readily calibrated using drug concentration data. The intrinsic tumor growth rate *r* was removed from consideration, as it can be readily measured from control growth experiments. We also removed the maximum rate of tumor kill by pembrolizumab *d* and the percent target occupancy in the TME that results in 50% kill by pembrolizumab *TO*_50_, as these values could conceivably be estimated through in vitro cell kill assays. The local sensitivity of the remaining parameters was then assessed (Figure S2). The four most sensitive parameters, in order, for fitting both the simulated %TO in the TME data and the TGI data (pembrolizumab administered on average every 3.5 days at a dose of 10 mg/kg) are: *k*_*synt*_, *K*_*x*_, *k*_1*T*_, *k*_*onT*_. We chose *k*_*synt*_ and *k*_*onT*_ for our analyses, as these parameters are unlikely to be estimated using any available experimental techniques but have proven to be critical in preceding work that analyzed a structurally similar model (Kareva et al. 2023).
4. *Generate simulated data for the measurement of interest*. Here we assume that both parameters of interest are normally distributed with a mean equal to the calibrated value of the parameter in Table 1. To get the desired variability in the simulated data, we choose the standard deviation for *k*_*onT*_ to be one fifth of its mean value (resulting in *k*_*onT*_∼𝒩(0.01,4 × 10^−6^)) and the standard deviation for *k*_*synt*_ to be one twentieth of its mean value (resulting in *k*_*synt*_∼𝒩(14190,503390.25)). We generate *K* = 10 random samplings of these distributions and extract the model-predicted value of percent target occupancy in the TME at each day over a one-month period. That is, for *k* = 1, …,10 we compute 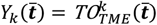, where 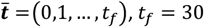 days. The resulting simulated percent target occupancy in the TME data is shown in Figure 4A.
5. *Practical identifiability using the data from “complete” simulated experiments*. Figure 4B,C shows that both *k*_*onT*_ and *k*_*synt*_ are practically identifiable when %TO in the TME is measured daily, given the parabolic shape of their profile likelihood curves.

**Figure 4.**
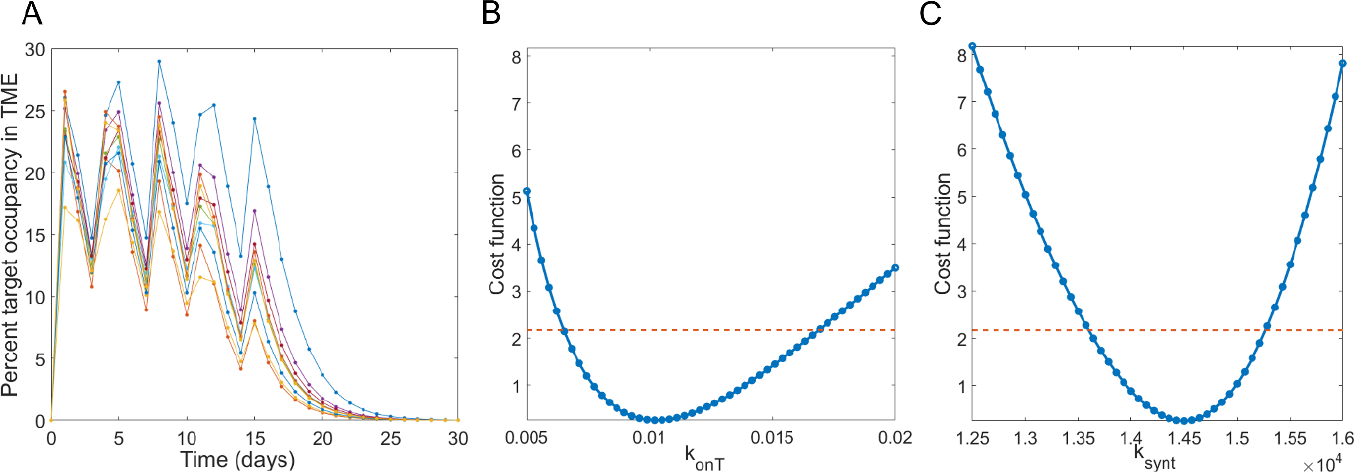
(A) shows 10 simulated datasets of percent target occupancy in the TME, generated using model parameters in Table 1 under assumptions *k*_*onT*_∼𝒩(0.01,4 × 10^−6^) and *k*_*synt*_∼𝒩(14190,503390.25). Profile likelihood curve for (B) *k*_*onT*_ and (C) *k*_*synt*_ when percent target occupancy in the TME is available every day (markers shown in (A)) indicate that both parameters are practically identifiable in this “complete” experimental scenario.

As we have verified that the parameter set ***p*** = (*k*_*onT*_, *k*_*synt*_) is practically identifiable when percent target occupancy in the TME data is collected every day for the duration of the experiment (complete data), we are ready to move to Step 6 of the method to search for the minimal number of data points that ensures practical identifiability of the parameters in ***p***.

### Insufficient Predictability With a 1-Day Experimental Protocol

We first sought to determine if the identifiability of ***p*** = (*k*_*onT*_, *k*_*synt*_) can be ensured with only a single measurement of percent target occupancy in the TME. To achieve this goal, we computed the profile likelihood curves for both *k*_*onT*_ and *k*_*synt*_ and examined whether practical identifiability is ensured if we only had a single time point available from the simulated dataset. If a single measurement is sufficient, we can also determine if the timing of the measurement (that is, what day it is taken) matters for preserving identifiability. The results of this analysis are shown in Figure 5.

**Figure 5.**
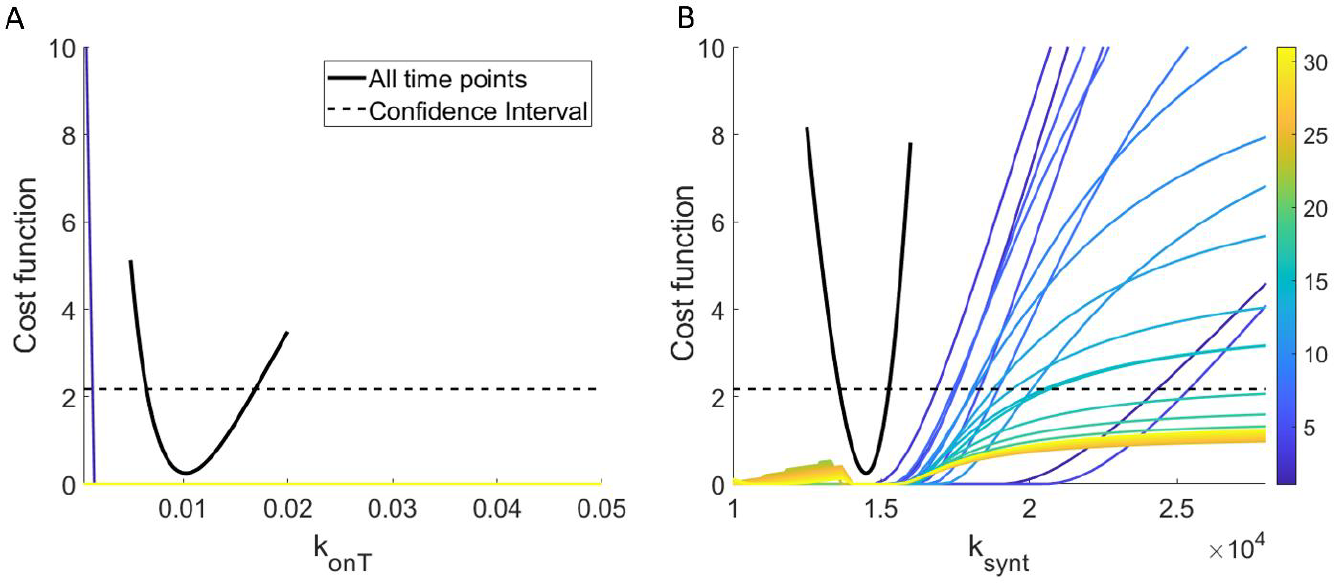
Profile likelihood curves for (A) *k*_*onT*_ and (B) *k*_*synt*_ using only a single experimental measurement of percent target occupancy in the TME. The color indicates which day (from day 1 to 30) the single experimental measurement was “collected”. The solid black curve is the profile likelihood curve for the specified parameter when all complete data (that is, daily measurements) are used, and the black dashed line is the corresponding 95% confidence threshold.

In all cases, we find that *k*_*onT*_ is not practically identifiable over the domain of interest. In fact, with one exception, the profiles appear flat (structurally non-identifiable) on a linear scale, though viewing the parameters on a log scale does reveal the profiles are not completely flat (see Figure S3A). If the experiment were conducted on day 1, *k*_*onT*_ would clearly be structurally identifiable, though it still not practically identifiable. In contrast, we find that generally *k*_*synt*_ is structurally identifiable, as the profiles are not flat. However, the parameter does not achieve practical identifiability independent of when the single measurement of target occupancy in the TME is taken. This is evident in Figure 5B as none of the profiles exceed the 95% confidence threshold as we both decrease and increase the parameter from its global minimum value.

These profiles lead us to conclude that a single experimental measurement of percent target occupancy in the TME is *insufficient* to confidently identify the values of either *k*_*onT*_ (rate of complex formation in the TME) or *k*_*synt*_ (rate of target synthesis in the TME). In other words, the practical identifiability of these parameters is lost (compared to the case of complete data) if we only have a single measurement of %TO in the TME, regardless of when that measurement is collected.

It is known that “parameter unidentifiability can lead to erroneous conclusions in the model inferences, predictions, and parameter estimates” (Eisenberg & Jain 2017). For this reason, we next explored the consequences of parameter non-identifiability on the proposed model’s predictive abilities. We do this by comparing the projected tumor volume over a parameter’s 95% confidence interval to the actual tumor volume in the experimental data to which the model was calibrated. These parameter sets are found by considering each profile likelihood curve generated from the experimental collection day of interest and identifying each value of the profiled parameter that falls below its 95% confidence threshold (see Figure S3A, C). For each such parameter value, pairing it with the corresponding best-fit value of the non-profiled parameter (see Figure S3B, D) forms what we call a “plausible parameter set” for the model, given the available data. We do note that other methods have been proposed for identifying the prediction confidence interval, including the prediction profile likelihood approach (Kreutz et al. 2012).

As shown in Figure 6A, when percent target occupancy in the TME is collected early (day 1), the range of predicted tumor volumes over its plausible parameter set is very large: the tumor is predicted to be anywhere from eradicated to more than 34 times the initial volume by day 30. Although the experimental data are all contained within these bounds, the model essentially has no predictive abilities if our single percent target occupancy in the TME data point is collected early, as the model cannot even infer whether the tumor volume decreases or increases due to treatment.

**Figure 6.**
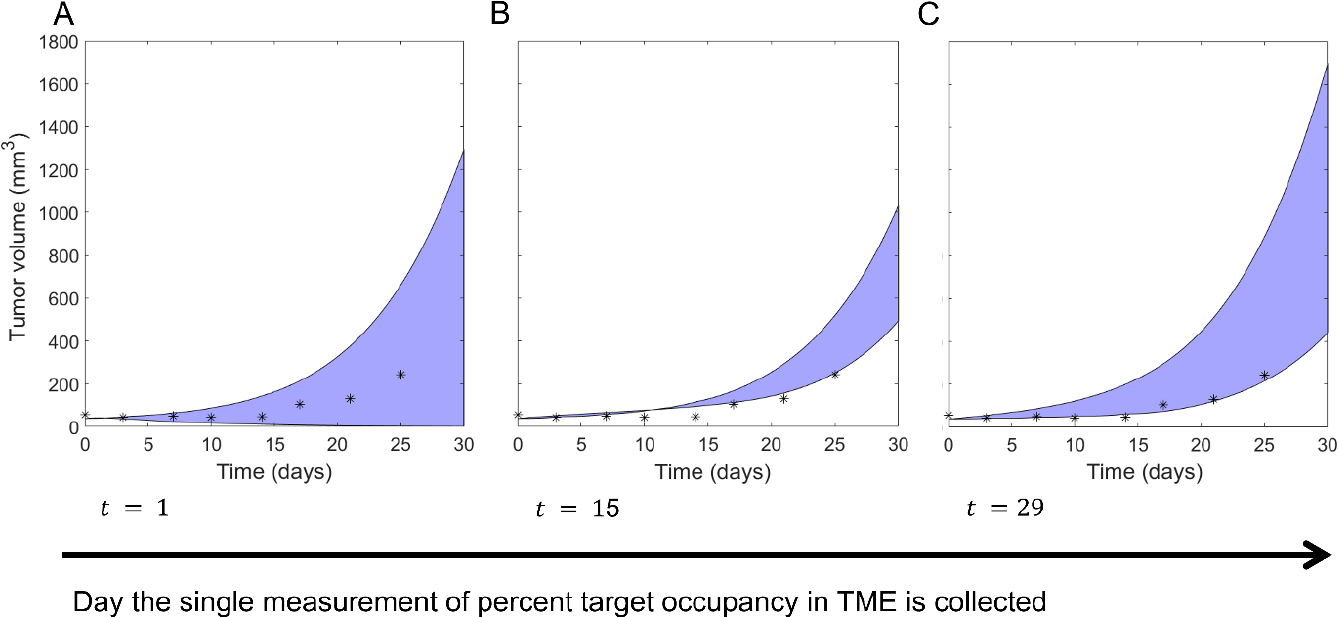
Shaded blue region shows the range for the predicted tumor volume over the plausible parameter sets when percent target occupancy in the TME is only measured at (A) the beginning of week 1 (day 1), (B) the beginning of week 3 (day 15), and (C) the beginning of week 5 (day 29). The true experimental measurement for tumor volume from (Agency 2015) is indicated with *s.

If instead percent target occupancy in the TME were measured at an intermediate time (day 15), the range of predicted tumor volumes over the plausible parameter set narrows significantly, as shown in Figure 6B. While a narrower predicted range is desirable, most of the experimental data does not lie within the model-predicted range for tumor volume. Thus, collecting percent target occupancy in the TME at an intermediate time point is also problematic. The best-case scenario occurs when the single measurement is collected near the end of the one-month period (Figure 6C). In this case, all plausible parameter sets predict that the tumor volume increases during and after treatment, and the experimental data is largely contained within the predicted range. That said, the tumor size at the end of the month is predicted to be anywhere from 441 – 1700 mm^3^. This large variation still highlights the problematic nature of using the model to predict treatment response when only a single measurement of percent target occupancy in the TME is available to calibrate model parameters.

In conclusion, we find that collecting a single measurement of percent target occupancy in the TME is insufficient for calibrating the model described in System (2) that bridges PK and PD components using a TME compartment. If we are in a situation where only a single measurement can be obtained, the experimental design recommendation is to collect that measurement as late in the month as possible.

However, this experimental design is far from ideal, as the unidentifiable parameters greatly limit confidence in the model’s predictions.

### Non-Robust Predictability over a Subset of 2-Day Experimental Protocols

We next sought to determine if the identifiability of ***p*** = (*k*_*onT*_, *k*_*synt*_) can be ensured using two measurements of percent target occupancy in the TME. Thus, we computed the profile likelihood curves for *k*_*onT*_ and *k*_*synt*_ when experimental data is collected at days (*t*_1_, *t*_2_), 0 < *t*_1_ < *t*_2_ ≤ 30. The resulting 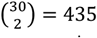 profile likelihood curves for each parameter are shown in Figure S4. We find that the structural non-identifiability issues that *k*_*onT*_ had using only a single measurement have been resolved, as none of the profiles are flat over their domain (Figure S4A). Further, it appears that some profiles for both *k*_*onT*_ and *k*_*synt*_ cross over the 95% confidence threshold, suggesting that some experiments that collect two measurements of percent target occupancy in the TME ensure practical identifiability.

To further investigate this, we classified all possible 2-day protocols for collecting %TO in TME at days (*t*_1_, *t*_2_) by whether they result in both, one of, or none of *k*_*onT*_ and *k*_*synt*_ being practically identifiable (Figure 7A). We find that only six of the 435 possible 2-day protocols result in both parameters being practically identifiable: (*t*_1_, *t*_2_) = (1,3), (3,4), (4,6), (4,7), (4,10), (4,13). The profiles for *k*_*onT*_ and *k*_*synt*_ corresponding to these 2-day protocols are shown in Figure 7B,C. Those profiles all have global minimum values that are very close to the true best-fit value of the parameter if complete experimental data were used (that is, if the percent target occupancy in the TME were collected daily). Thus, if these data were collected using one of the six identifiable 2-day protocols, we can confidently estimate the value of parameters ***p*** = (*k*_*onT*_, *k*_*synt*_).

**Figure 7.**
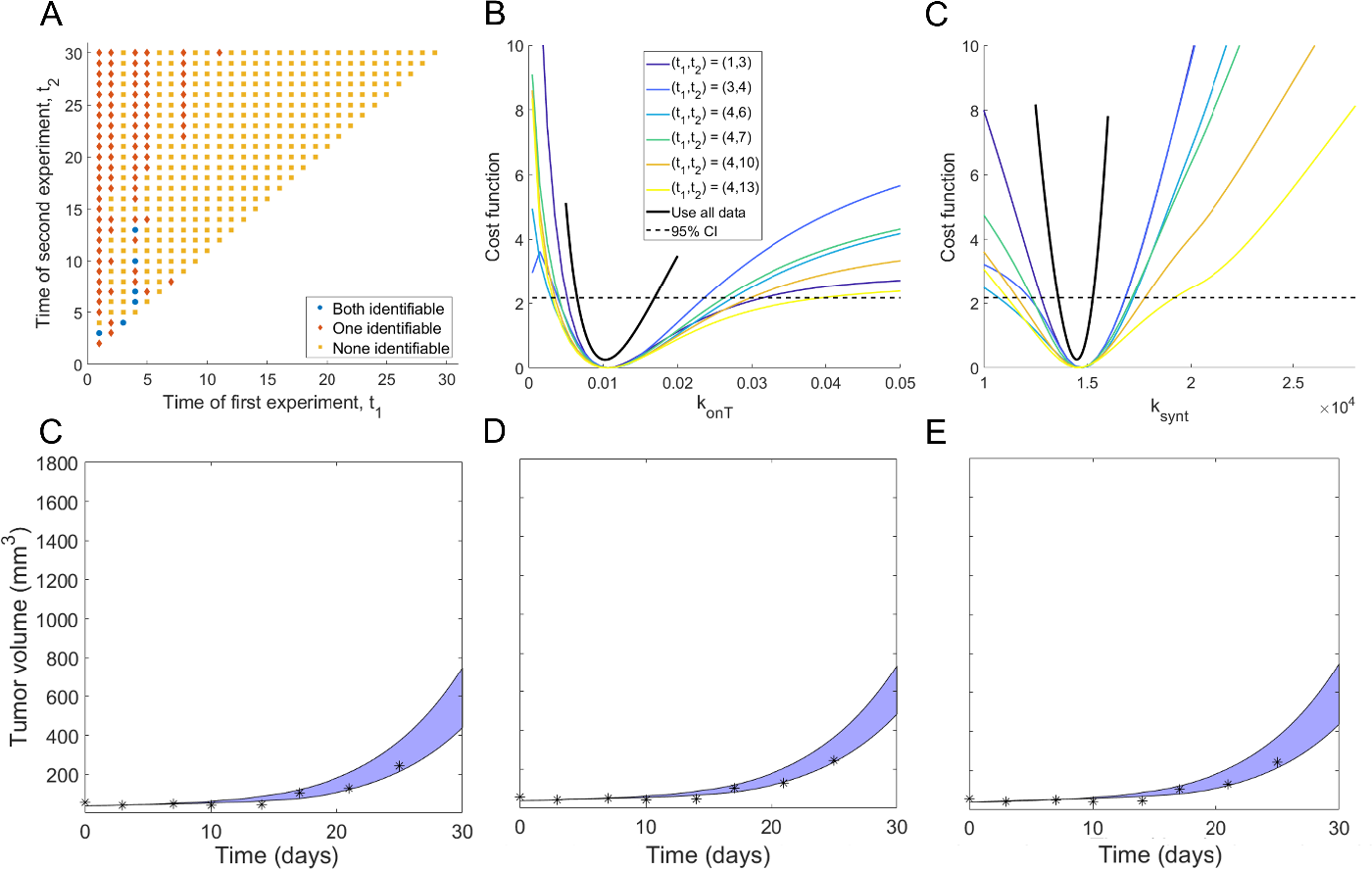
(A) Classification of all possible 2-day protocols (*t*_1_, *t*_2_) by whether they result in both (blue circles), one (red diamonds) or none (orange squares) of the parameters being practically identifiable. The profile likelihood curves using all 2-day protocols that result in practically identifiable parameters are shown in (B) for *k*_*onT*_ and (C) for *k*_*synt*_. (D)-(F) Blue region shows the range for the predicted tumor volume over the plausible parameter sets for three of the six 2-day experimental protocols that correspond to practically identifiable parameters. (*t*_1_, *t*_2_) = (1,3) in (D), (3,4) in (E) and (4,6) in (F). Note the *y*-scale is the same as in Figure 6 to clearly highlight the predictive differences when using protocols that do not result in parameter identifiable, versus protocols that do.

We next explore the validity of model TGI predictions using the 2-day experimental protocols which resulted in practically identifiable parameters. As with the 1-day protocols, we do this by simulating the predicted tumor volume over the “plausible parameter sets”. However, this time we restrict ourselves to considering the 2-day protocols that correspond to both parameters being practically identifiable. As shown in Figure 7(D)-(F), the predicted tumor region is highly constrained, and most of the experimental data lies within this model-predicted range. This demonstrates that the model can well predict the experimental tumor volume measurements, provided one of the identified 2-day protocols was used to collect percent target occupancy in the TME.

Notably, even though our approach has identified six 2-day experimental protocols that achieve our goal of preserving the identifiability of *k*_*onT*_ and *k*_*synt*_ (and thus result in confident model predictions of TGI), we are disinclined to recommend designing an experiment based around them as the identified protocols are not robust. To detail, if we perturb any of the recommended 2-day protocols by a single day (that is, if we consider (*t*_1_ ± 1, *t*_2_), (*t*_1_, *t*_2_ ± 1) for any of the recommended protocols), only two of the perturbed protocols ensure the practical identifiability of both parameters. This can be clearly seen in Figure 7A, which shows that experiments with both parameters classified as practically identifiable (blue circles) generally have neighboring points for which either none (orange squares) or one (red diamonds) of the parameters are identifiable. Thus, even though there do exist protocols that collect percent target occupancy in the TME at two days that can well-inform model parametrization, we recommend against this experimental design given its sensitivity to the precise timing for when the data must be collected.

### Robust Identifiability over a Subset of 3-Day Experimental Protocols

Given the lack of robustness of the discovered 2-day protocols, we next sought to determine if identifiability of ***p*** = (*k*_*onT*_, *k*_*synt*_) can be ensured using three measurements of percent target occupancy in the TME. There are 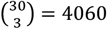 such 3-day protocols. Generating the profile likelihood for each of these protocols becomes computationally expensive, even in our scenario where only two parameters are being profiled. Thus, rather than generate profile likelihood curves for all 4060 protocols, we instead select a random third of the protocols for our analysis and assume that this sampling is sufficient to understand the entirety of the 3-day protocol space. The sufficiency of analyzing only a fraction of the possible protocols is verified below (Figure 8E,F). Removing any non-unique random samplings, this resulted in the consideration of 1126 3-day protocols. The profile likelihood curves corresponding to each of these 3-day protocols are found in Figure S5. We observe that a number of these protocols appear practically identifiable.

**Figure 8.**
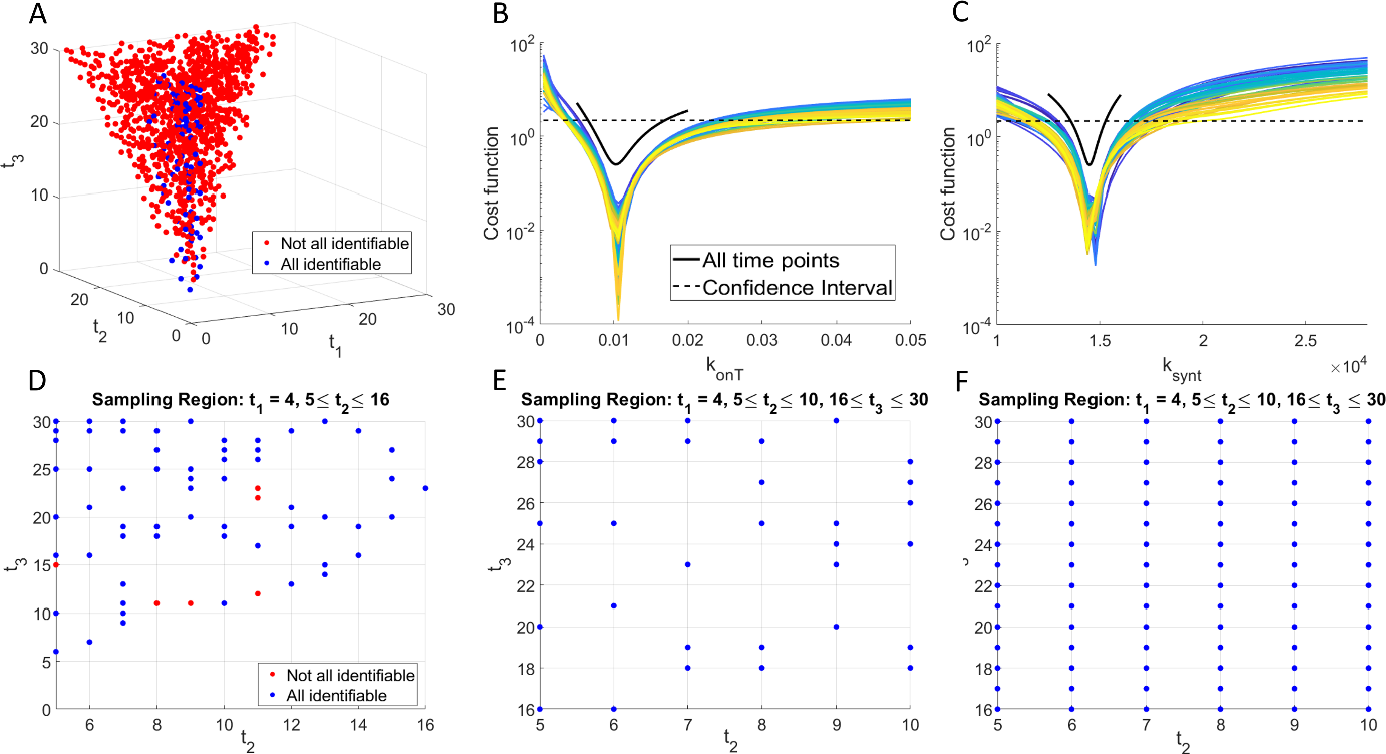
(A) Classification of a random sampling of 3-day protocols (*t*_1_, *t*_2_, *t*_3_) by whether they result in both parameters being practically identifiable (blue circles) or not (red circles). The profile likelihood curves from the randomly sampled 3-day protocols that result in practically identifiable parameters are shown in (B) for *k*_*onT*_ and (C) for *k*_*synt*_. (D) Projection of plot in (A) with *t*_1_ = 4 and *t*_2_ ≥ 5. (E) Further narrowing the protocol space to the region *P* defined in eqn. (3) results in all sampled protocols contained in *P* being practically identifiable. (F) Testing all protocols in *P*, not just randomly sampled ones, confirms that any protocol in *P* results in both parameters being practically identifiable.

To investigate further, we classified all possible 3-day protocols (*t*_1_, *t*_2_, *t*_3_) by whether they result in both *k*_*onT*_ and *k*_*synt*_ being practically identifiable (Figure 8A). We find that 80 of the 1126 3-day protocols considered result in both model parameters being practically identifiable. The profiles for *k*_*onT*_ and *k*_*synt*_ corresponding to these 3-day protocols are shown in Figure 8B,C. Those profiles all have global minimum values that are very close to the true best-fit value of the parameter if complete experimental data were used (that is, if the percent target occupancy in the TME were collected daily). Thus, if these data were collected using any of the 80 identifiable 3-day protocols, we can confidently estimate the value of parameters ***p*** = (*k*_*onT*_, *k*_*synt*_).

An analysis of the data in Figure 8A shows that 73.75% of the 3-day protocols that correspond to practically identifiable parameters (59 of 80) collect the first measurement of percent target occupancy in the TME at day *t*_1_ = 4. Further, the second measurement must be taken after the first, but by day *t*_2_ = 16 if we want to ensure parameter identifiability. Fixing *t*_1_ = 4 in protocol space allows us to project our results into a two-dimension plot in *t*_2_ − *t*_3_ space (Figure 8D, where we also impose 5 ≤ *t*_2_ ≤ 16). This region contains 65 of our tested 3-day protocols, 90.77% of which are practically identifiable. Thus, an experimental protocol with *t*_1_ = 4, 5 ≤ *t*_2_ ≤ 16, *t*_3_ > *t*_2_ is very likely to result in identifiability of both parameters, but it is not guaranteed to do so. In Figure 8E, we show that if we further narrow down protocol space to

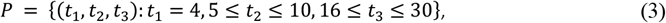

then every 3-day protocol we had randomly sampled in the region *P* results in practically identifiable parameters. We note that there are many practically identifiable protocols outside of *P*, as this region only contains 40% of all the practically identifiable 3-day protocols we tested. That said, we find using a protocol in *P* to be highly desirable, as the experimental design is robust to small changes in the day the measurements are collected. We confirm this robustness in Figure 8F, where we tested the identifiability associated with every protocol in *P*, not just the ones that had been randomly sampled during our analysis. As predicted from the random sampling, every protocol in *P* corresponds to both model parameters being practically identifiable. This also confirms the sufficiency of using a random sampling of possible protocols in the proposed minimal experimental design methodology.

In conclusion, our identifiability analysis has led to a robust, minimally sufficient, experimental design recommendation. We suggest an experimental protocol where percent target occupancy in the TME is collected three times over a one-month period. The first sample should be collected on day *t*_1_ = 4. The second sample can be taken anywhere from day 5 to day 10, 5 ≤ *t*_2_ ≤ 10. The third sample should be taken on day 16 or later, 16 ≤ *t*_3_ ≤ 30. Even though approximately 60% of 3-day protocols that result in practically identifiable parameters lie outside the recommended region, we have shown that this experimental design guarantees that every choice made using these specifications results in practically identifiable parameters. Using any experimental design away from the boundary of this region is thus predicted to robustly provide sufficient data for confidently estimating the values of the parameters ***p*** = (*k*_*onT*_, *k*_*synt*_).

### Sensitivity of Experimental Design Recommendation to Methodological Assumptions

A number of decisions must be made to implement the proposed minimal experimental design algorithm. Beyond the standard decisions that go into model development (building the right-sized model for the problem at hand, given the available data), a subset of model parameters must be chosen for further study using a combination of experimental knowledge and sensitivity results (see Figure S2).

Once these parameters have been selected, the modeler has another decision to make: from what underlying distribution should the parameter values be sampled to generate the simulated data needed at the fourth step (see Figure 3) of the workflow?

In the results presented thus far, we assumed that the parameters are normally distributed. The mean of each distribution is the best-fit value of the parameter when the model is fit to TGI data (10 mg/kg of pembrolizumab administered on average every 3.5 days). The standard deviations were selected to strike a balance between being too “optimistic” (i.e., having such low variability that it is unlikely to represent real data) and being too “pessimistic” (i.e., having such high variability that the experimental data are too noisy to be informative). In Table 2, we explore what happens if we change the assumptions placed on the parameter distributions. We particularly ask how the experimental design recommendation changes if: 1) parameters are lognormally distributed instead of normally distributed (keeping the mean and standard deviation fixed), 2) parameters remain normally distributed about the same mean but the standard deviation is decreased by 25%, and 3) parameters remain normally distributed about the same mean but the standard deviation is increased by 25%.

**Table 2.** Robustness of Experimental Design Recommendation to Parameter Distributions. Number in table is the percent of protocols for which parameters are practically identifiable.

| **Protocol**<br>**Distribution** | **Measure %TO**<br>**in TME at 1**<br>**time point** | **Measure %TO in TME**<br>**at 2 time points** | **Measure %TO**<br>**in TME at 3**<br>**time points** |
| --- | --- | --- | --- |
| **Normal (default)** | 0% | 1.38%<br>Identifiable Protocols:<br>(1,3), (3,4), (4,6), (4,7), (4,10), (4,13) | 7.10% |
| **Lognormal**<br>**($\mu$, $\sigma$ unchanged)** | 0% | 1.61%<br>Identifiable Protocols:<br>(1,3), (3,4), (4,7), (4,9), (4,10), (4,13) | 7.02% |
| **Normal (0.75σ)** | 0% | 2.99%<br>Identifiable Protocols: (1,3),<br>(1,6), (1,7), (1,9), (1,10), (1,13), (1,14),<br>(3,4), (4,6), (4,7), (4,10), (4,13), (4,14) | 16.16% |
| **Normal (1.25σ)** | 0% | 0.02%<br>Identifiable Protocol: (4,7) | 3.46% |

One conclusion is consistent across all cases considered: measuring %TO in the TME at a single time point is not a sufficient experimental design, as it never results in practically identifiable parameters.

Focusing next on changing the shape of the distribution, we find that shifting from a normal to a lognormal distribution (with the same mean and standard deviation) has minimal impact. Table 2 shows that a very similar set of 2-day protocols results in practically identifiable parameters, and a very similar percentage of 3-day protocols result in practically identifiable parameters. Figure S6A shows the classification of all tested 3-day protocols and Figure S6B demonstrates that all protocols in the experimentally-recommended region *P* identified using a normal distribution result in practically identifiable parameters when the simulated data is generated from a lognormal distribution. Thus, the experimental design recommendation made using normally distributed parameters holds if we change the distribution to lognormal.

We next consider the impact of keeping the parameters normally distributed about the same mean but changing the standard deviation. The experimental design recommendation is influenced by the standard deviation in the following way: the smaller the standard deviation, the larger the set of experimental protocols that result in practically identifiable parameters. To detail, 2.99% of 2-day protocols result in practically identifiable parameters when the standard deviation is 25% smaller than the default value. That number drops to 1.38% at the default standard deviation, and 0.02% when the standard deviation is 25% larger than the default value. The decrease in the number of identifiable protocols as a function of standard deviation is to be expected, as more noisy data should necessitate the collection of more data (Harshe et al. 2023). Table 2 shows that every protocol that resulted in practically identifiable parameters using the default value of σ also results in practically identifiable parameters at 0.75σ. Thus, a smaller standard deviation means more experimental designs are allowable, but it also preserves the recommendations made using (somewhat) more noisy data.

Looking at the randomly sampled 3-day protocols, we see the same qualitative trend as a function of standard deviation as with the 2-day protocols. 16.16% of protocols result in practically identifiable parameters at 0.75σ (see Figure S6C), compared to 7.10% at σ and 3.46% at 1.25σ (see Figure S6E). All protocols in region *P* identified using the default value of σ still result in practically identifiable parameters using 0.75σ (Figure S6D). At 1.25σ, only a subset of the protocols in region *P* result in practically identifiable parameters. In particular, collecting the second sample on day 5 or 8 is no longer a recommended experimental design. Thus, a larger standard deviation imposes more restrictions on when data should be collected.

## DISCUSSION

As mathematical models continue to be used for real time interventions, it becomes increasingly important to design experiments that collect the right data, at the right time, to maximize the model’s predictive power. In this work, we propose an approach for using identifiability analysis to design a minimally sufficient set of experiments. The approach involves generating simulated data from the model, and then identifying a set of parameters that would be practically identifiable in the ideal scenario of “complete” experimental data (i.e., conducting an experiment daily). Then, we work backwards, seeking to find the minimal number of data points that must be collected to ensure the practical identifiability of selected model parameters. We applied the proposed minimal experimental design technique to a site-of-action PKPD model of tumor growth in response to treatment with pembrolizumab that explicitly accounts for drug-target binding in the TME. We were particularly interested in identifying two very specific quantities that are difficult to measure experimentally but that significantly affect model dynamics – target synthesis in the TME and apparent drug-target affinity in the TME.

Applying our minimally sufficient experimental design algorithm led to the conclusion that, even if all other parameters are estimated using other methods and experiments, no single measurement of percent TO in the TME can result in parameter practical identifiability. Consequently, the model cannot be confidently parameterized, and the resulting predictions about tumor response to treatment cannot be trusted. We did identify a small number of 2-day experimental protocols for collecting %TO in the TME that ensure parameter practical identifiability. However, these protocols are not robust to small perturbations in when the data is collected. Further, if the experimental data were somewhat noisier (25% more) than assumed, only one of the identified 2-day protocols result in practically identifiable parameters. It took considering 3-day protocols to identify a robust set of experimental designs that ensure parameter identifiability: the first measurement of %TO in the TME must be collected on day 4, the second can be collected any time between days 5 and 10 (inclusive), and the third taken any time from day 16 onward. Any such set of three measurements is sufficiently informative for the selected parameters to be practically identifiable, which lends strong confidence to predictions made by the model. This recommendation is fairly robust to the underlying assumptions about the simulated data: the recommendation holds if the data is lognormally distributed instead of normally distributed and if the data is normally distributed but 25% less noisy than assumed. If the data is normally distributed and 25% noisier than assumed, a subset of the recommended protocols will result in practically identifiable parameters.

While the sampling schedule recommended here is particular to the model and experimental system under consideration, the proposed minimally sufficient experimental design methodology can be applied broadly to other biological systems. The method is best applied, however, to design an experiment that yields important data for model parametrization but is challenging/invasive/expensive to perform. For instance, there is no need to optimize the experimental design for measuring percent target occupancy in the TME if sufficient dynamics can be inferred from the correlation between plasma PK and tumor burden reduction. Similarly, since collecting volumetric data in mice is straightforward and non-invasive, there is no need to design a minimally sufficient experiment for this task. Conversely, there are times when reliable understanding of TO dynamics in the TME is critical, as would be the case for bispecific T cell engagers (BiTes), whose efficacy is contingent on the drug binding to both targets explicitly in the TME (Vafa & Trinklein 2020). Additionally, assessing levels of target engagement can be very important for addressing safety considerations, both for BiTEs (Saber et al. 2017), and for broader classes of drugs, such as immune cell agonists (Muller et al. 2009). For these cases, the additional investment of resources in data collection even for an invasive procedure may prove indispensable. The proposed methodology enables collecting the minimal amount of data sufficient to inform an associated mathematical model.

A key to successful implementation of the proposed minimally sufficient experimental design methodology is determining the set of parameters for analysis. The set must be small enough to ensure that all parameters are practically identifiable under the “ideal” experimental conditions; for instance, if the data of interest could be collected daily over the entire course of the experiment. Any parameters not in the analysis set must be fixed, thus posing the challenge of how to approximate the value of such fixed parameters. Fortunately, for PKPD models, some common approaches exist for parameter estimation. In the proposed example, for instance, PK parameters can be estimated using standard software, such as Monolix or WinNonLin; *k*_*on*_ and *k*_*off*_ can be assessed using Biacore (Malmborg & Borrebaeck 1995; Jason-Moller et al. 2006) as a first step and then refined using in vitro assays to further capture the expected PK-TO relationships. Target clearance rates can be estimated using internalization assays, and normal synthesis rates can be calculated from expected normal steady-state levels of the target. With such careful preparatory work, this technique enables estimation of the few remaining elusive parameters that may be essential to a model’s predictive value.

The benefits of the proposed experimental design methodology are best realized when experimentalists and modelers are working in close collaboration. Such a partnership ensures alignment of key measurable aspects of the mechanism of interest, while ensuring that modelers have a clear understanding of what experiments are feasible. Constraints on what experiments can be conducted include experiment duration, the possible number of measurements that can be collected, and any constraints on when the samples can be collected (due to costs, implementation issues, animal welfare, etc.). While optimizing time point collection does not answer all experimental design questions – for instance, it does not answer questions related to selecting sample size – the proposed methodology can be used in combination with other optimal experimental design approaches to maximize the utility of collected data.

More broadly, this work reinforces the idea that there is no “correct” model in absolute; instead, there are data for the model and model for the data. Indeed, when we talk about fit-for-purpose models, we typically talk about models that are just complex enough to answer the question of interest. This work arguably adds another dimension to fit-for-purpose modeling, where fit-for-purpose data are collected to inform the model to then enable it to answer the underlying question. Therefore, the structure of a model, even for the same question, can conceivably be different but equally useful depending on the data available to identifiably parametrize it. Once a research team has identified a motivating question and built and parameterized a fit-for-purpose model for the question of interest, the proposed experimental design framework can be a powerful tool for identifying the minimal amount of experimental data required to maximize the model’s predictive power.

## Data availability

The data analyzed in this study were accessed from previously published experimental literature. Citations to all sources of data are included in the manuscript.

## Code availability

The underlying code, and the training/validation datasets, for this study are available at Github and can be accessed via this link: https://github.com/jgevertz/minimal_experimental_design.

## AUTHOR CONTRIBUTIONS

JLG and IK jointly designed the research, analyzed the data, and wrote the manuscript. JLG performed the research. All authors read and approved the final manuscript.

### ACKNOWLEDGEMENTS

JLG acknowledges use of the ELSA high-performance computing cluster at The College of New Jersey for conducting the research reported in this paper. This cluster is funded in part by the National Science Foundation under grant numbers OAC-1826915 and OAC-1828163.

## COMPETING INTERESTS

Author IK is an employee of EMD Serono, the US business of Merck KGaA but declares no non-financial competing interests. The research was conducted during JLG’s sabbatical with EMD Serono in the Quantitative Pharmacology Department but declares no non-financial competing interests.

## Supplementary Information

**Figure S1.**
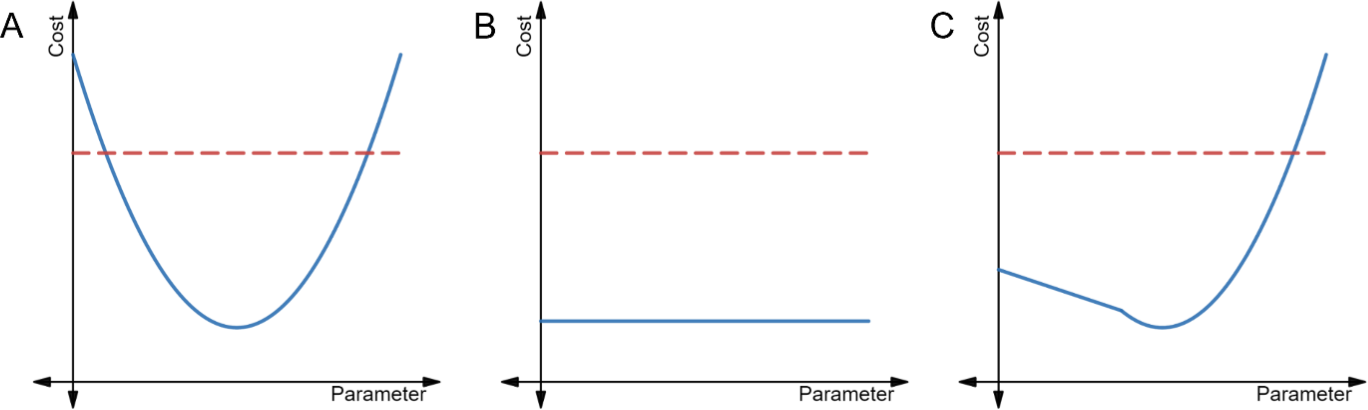
Illustration of the profile likelihood method. Profile likelihood curves (blue) of (A) a practically identifiable parameter, (B) a structurally non-identifiable parameter, and (C) a structurally identifiable but practically non-identifiable parameter. Thresholds for the 95% confidence intervals are indicated with red dashed lines. The figure is adapted from (Craig et al., 2023).

**Figure S2.**
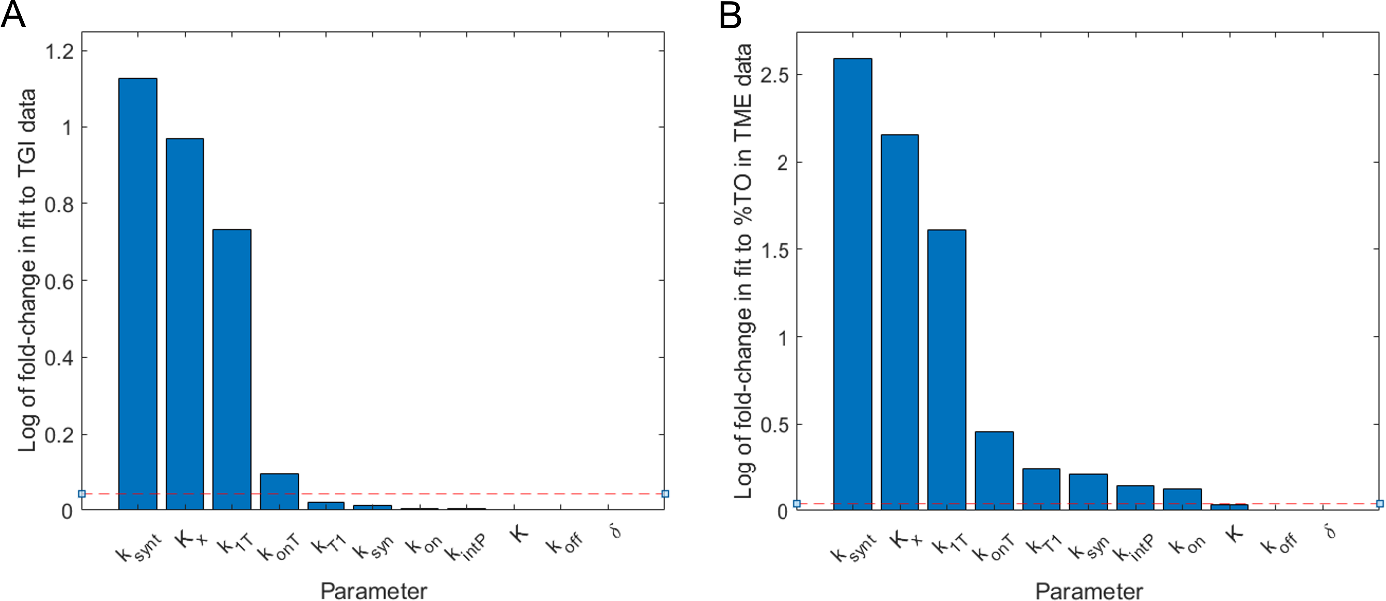
Local sensitivity analysis of a subset of model parameters. Each parameter was individually varied by ±10%, and the maximum of the log of the fold-change in model fit (relative to the best fit parameter set) was computed. (A) Sensitivity of fit to TGI data. (B) Sensitivity of fit to %TO in the TME data. The dashed line indicates the fold-change in the cost function that is equal to the fold-change in the parameter value.

**Table S1.** Computational resources and costs associated with various steps of the proposed minimal experimental design workflow. Note that, unless otherwise noted, all profiles here were generated by using 50 points uniformly distributed over each parameter’s domain.

| Code | Where/how ran | Run time | How to speed up run time |
| --- | --- | --- | --- |
| Fit model to simulated %TO in TME data and generate parameter profiles using all simulated data | Locally, not parallelized | @21min | N/A (Only used 12 points per profile) |
| Generate parameter profiles using every possible 1-day protocol (30 such protocols) | Locally, and parallelized to run over 4 "tasks" | @4.5h | 1. Use less than 50 points per profile<br>2. Use only a subset of 1-day protocols (ex: test every other day)<br>3. Parallelize over more "tasks" |
| Generate parameter profiles using every possible 2-day protocol (435 such protocols) | High performance cluster, parallelized over 63 "tasks" | @6.8h | 1. Use less than 50 points per profile<br>2. Use only a subset of 2-day protocols (a random N%) |
| Generate parameter profiles using a random 1/3 of all possible 3-day protocols and removing non-unique samples (tested 1126 of 4060 protocols) | High performance cluster, parallelized over 63 "tasks" | @16.7h | 1. Use less than 50 points per profile<br>2. Use less than 33% of all 3-day protocols |

**Figure S3.**
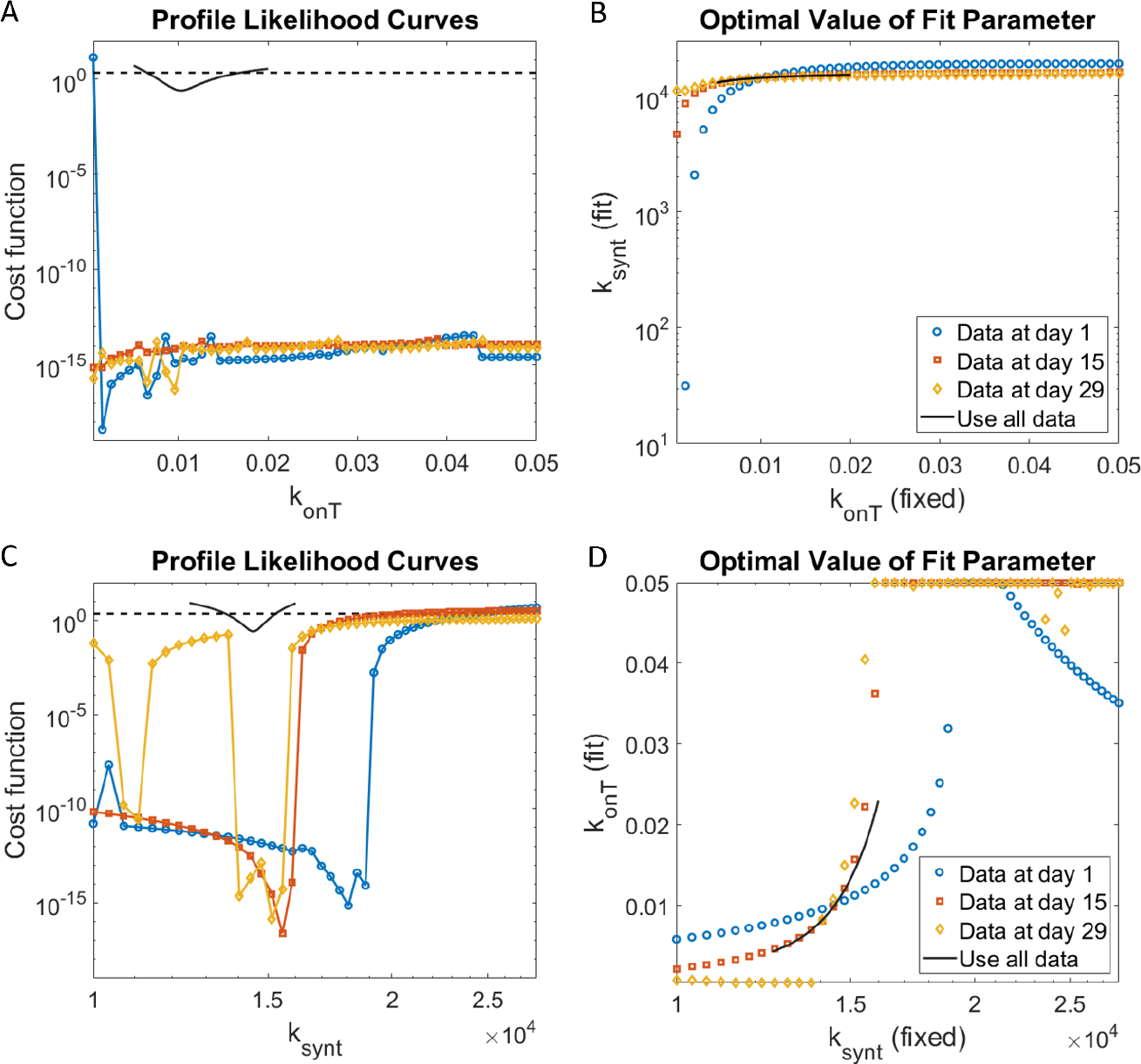
Profile likelihood curves for (A) *k*_*onT*_ and (C) *k*_*synt*_ when percent target occupancy in the TME is collected only on day 1 (blue circle), day 15 (red square), or day 29 (orange diamond). (B) shows the best-fit value of *k*_*synt*_ for each fixed value of *k*_*onT*_. (D) shows the best-fit value of *k*_*onT*_ for each fixed value of *k*_*synt*_. Black solid lines represent results using complete data (daily measurement of %TO in TME), and the black dashed line is the 95% confidence threshold for the profiled parameter using all complete data. Any parameter whose cost falls below the 95% confidence threshold, paired with the corresponding best-fit value of the fit parameter, form a “plausible” parameter set that is considered in Figure 6. It is of note that we restrict 0.0005 ≤ *k*_*onT*_ ≤ 0.05 when performing parameter fitting.

**Figure S4.**
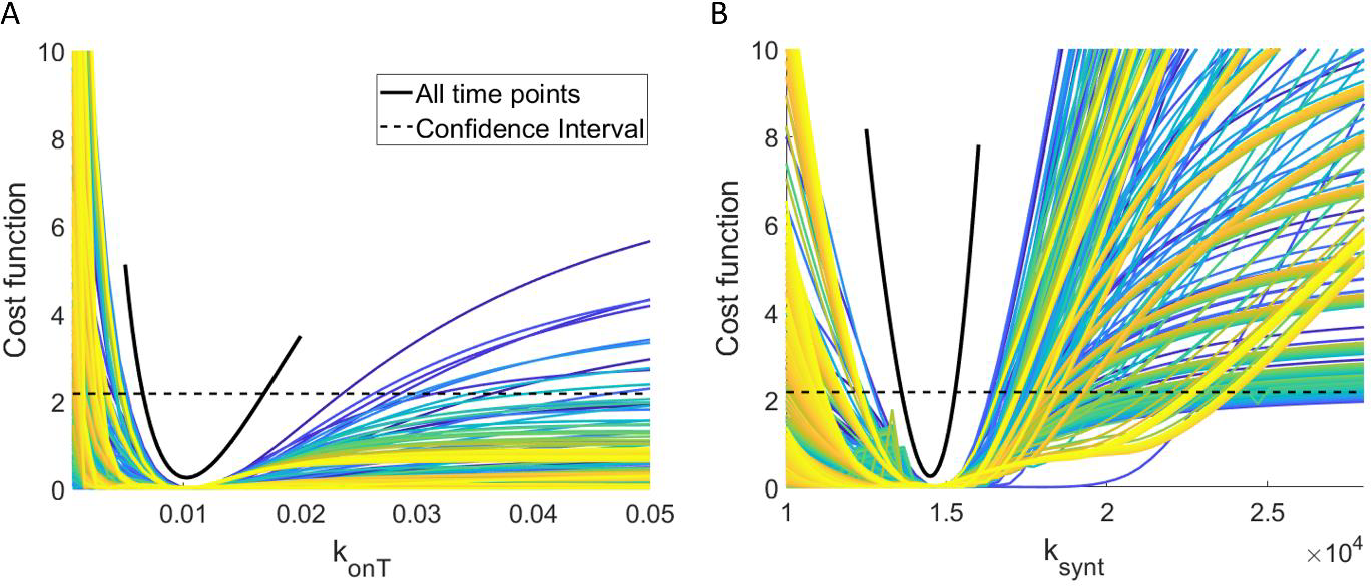
Profile likelihood curves for (A) *k*_*onT*_ and (B) *k*_*synt*_ for all 435 protocols that collect percent target occupancy in the TME at exactly two days. The color indicates the spacing between collection days, ranging from spacing the experiments by one day (blue) to 29 days (yellow). The solid black curve is the profile likelihood curve for the specified parameter when complete data is used, and the black dashed line is the corresponding 95% confidence threshold.

**Figure S5.**
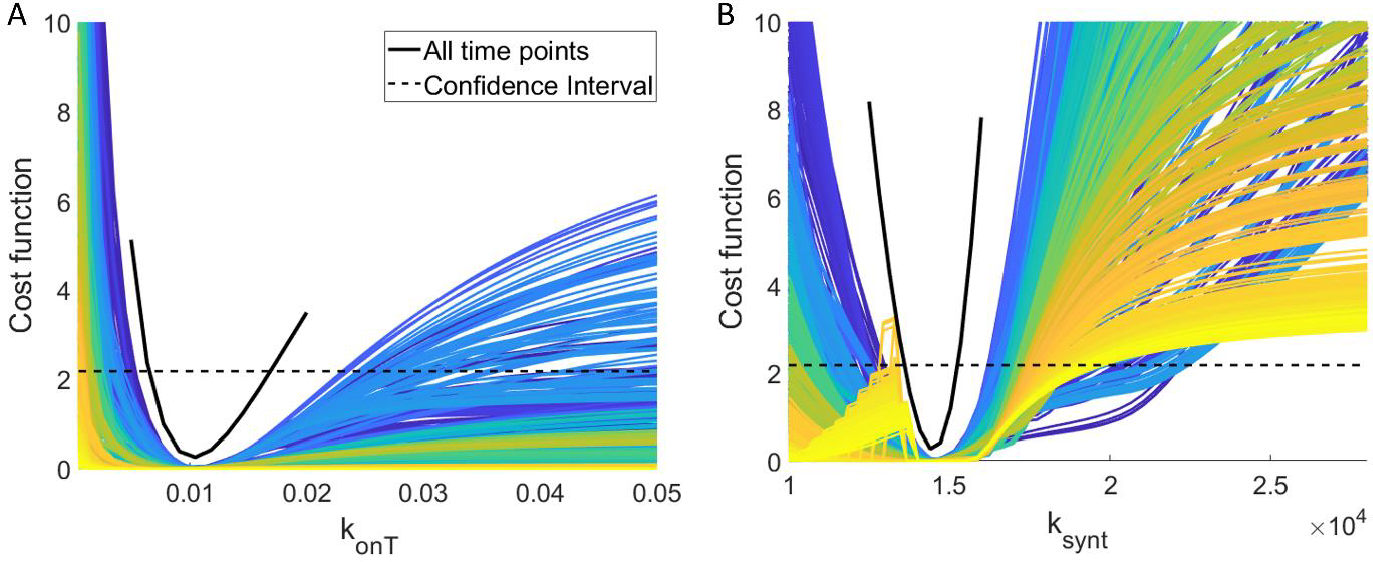
Profile likelihood curves for (A) *k*_*onT*_ and (B) *k*_*synt*_ for a random sampled set of 1126 protocols that collect percent target occupancy in the TME at exactly three days. The color indicates the spacing between the first two collection days, ranging from spacing the first two experiments by one day (blue) to 28 days (yellow). The solid black curve is the profile likelihood curve for the specified parameter when complete data are used, and the black dashed line is the corresponding 95% confidence threshold.

**Figure S6.**
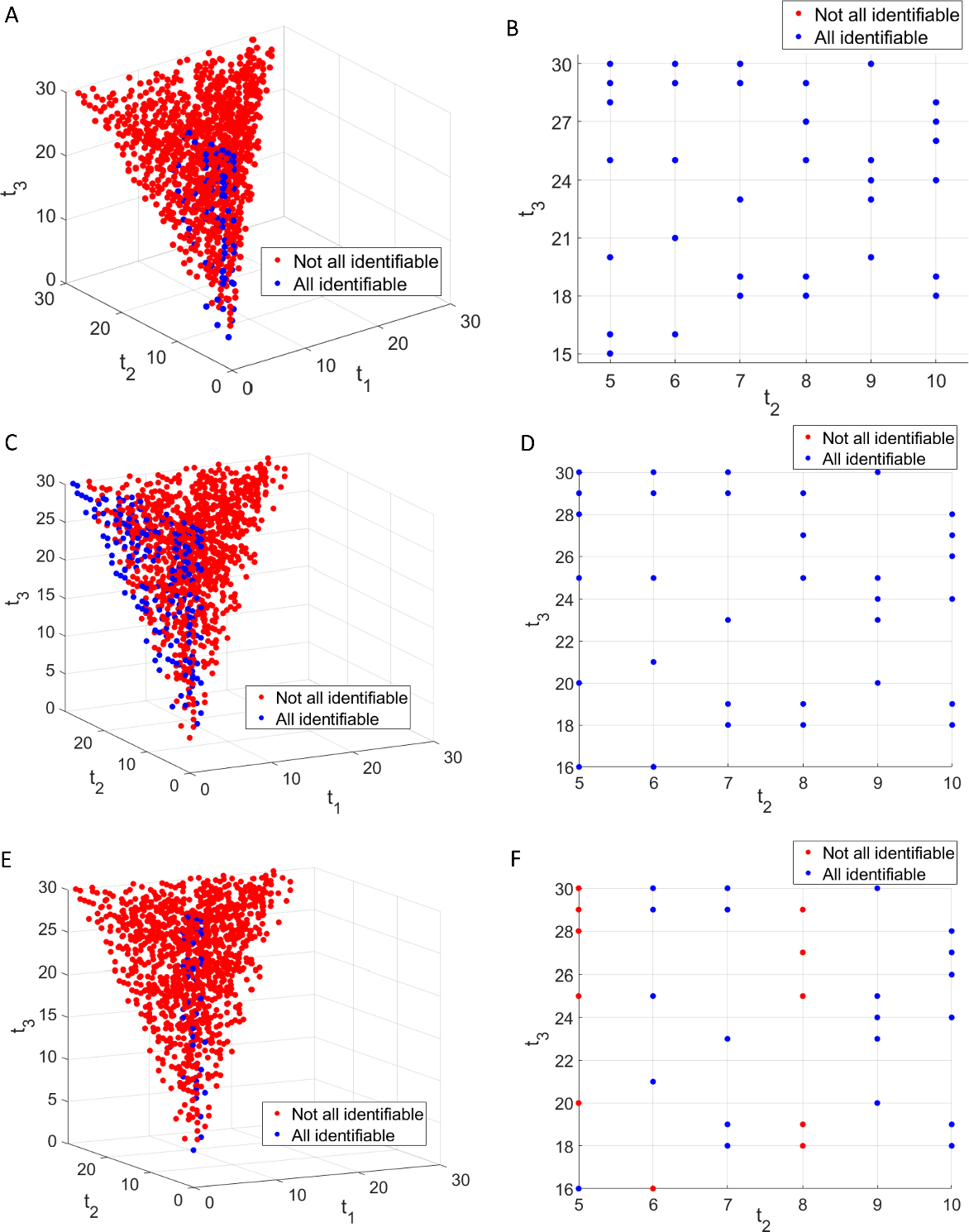
Exploring robustness of the experimental design recommendation to parameter distribution shape and standard deviation. Top row: using a lognormal distribution with same mean and standard deviation as the default normal distribution. Middle row: using a normal distribution with same mean as default normal distribution, but the standard deviation is 25% smaller. Top row: using a normal distribution with same mean as default normal distribution, but the standard deviation is 25% larger. First column: The classification of a random sampling of 3-day protocols (*t*_1_, *t*_2_, *t*_3_) by whether they result in both parameters being practically identifiable (blue circles) or not (red circles). Second column: Only shows the protocols found in the recommended protocol design region *P* (determined using the default value of σ).

## Notes

### Competing Interest Statement

The authors have declared no competing interest.

## References

Agency, E.M., 2015. Keytruda assessment report. Keytruda INN-pembrolizumab. Available at: https://www.ema.europa.eu/en/documents/assessment-report/keytruda-epar-public-assessment-report_en.pdf.

Beckman, R.A., Kareva, I. & Adler, F.R., 2020. How Should Cancer Models Be Constructed? Cancer Control, 27(1), p. 1073274820962008.

Betts, A. et al., 2019. A translational quantitative systems pharmacology model for CD3 bispecific molecules: application to quantify T cell-mediated tumor cell killing by P-cadherin LP DART®. The AAPS journal, 21, pp. 1–16.

Brady, R. & Enderling, H., 2019. Mathematical models of cancer: when to predict novel therapies, and when not to. Bulletin of mathematical biology, 81(10), pp. 3722–3731.

Buchwald, A.G. et al., 2021. Estimating the impact of statewide policies to reduce spread of severe acute respiratory syndrome coronavirus 2 in real time, Colorado, USA. Emerging Infectious Diseases, 27(9), p. 2312.

Cárdenas, S.D. et al., 2022. Model-informed experimental design recommendations for distinguishing intrinsic and acquired targeted therapeutic resistance in head and neck cancer. npj Systems Biology and Applications, 8(1), p. 32.

Cassidy, T., 2023. A continuation technique for maximum likelihood estimators in biological models. arXiv preprint arXiv:2303.09194.

Cho, H., Lewis, A.L. & Storey, K.M., 2020. Bayesian information-theoretic calibration of radiotherapy sensitivity parameters for informing effective scanning protocols in cancer. Journal of clinical medicine, 9(10), p. 3208.

Chudasama, V.L. et al., 2015. Simulations of site-specific target-mediated pharmacokinetic models for guiding the development of bispecific antibodies. Journal of pharmacokinetics and pharmacodynamics, 42(1), pp. 1–18.

Csilléry, K. et al., 2010. Approximate Bayesian computation (ABC) in practice. Trends in ecology & evolution, 25(7), pp. 410–418.

Dunlap, T. & Cao, Y., 2022. Physiological Considerations for Modeling in vivo Antibody-Target Interactions. Frontiers in Pharmacology, 13.

Eisenberg, M.C. & Jain, H.V., 2017. A confidence building exercise in data and identifiability: Modeling cancer chemotherapy as a case study. Journal of theoretical biology, 431, pp. 63–78.

Harshe, I., Enderling, H. & Brady-Nicholls, R., 2023. Predicting Patient-Specific Tumor Dynamics: How Many Measurements Are Necessary? Cancers, 15(5), p. 1368.

Hu, S., 2004. Optimal time points sampling in pathway modelling. In The 26th Annual International Conference of the IEEE Engineering in Medicine and Biology Society. IEEE, pp. 671–674.

Jason-Moller, L., Murphy, M. & Bruno, J., 2006. Overview of Biacore systems and their applications. Current protocols in protein science, 45(1), pp. 19–13.

Kareva, I. et al., 2023. Integrated model-based analysis utilizing co-expressed checkpoint inhibitor data to inform the recommended dose for expansion (RDE) of anti-TIGIT mAb M6223. Clinical Pharmacology & Therapeutics ASCPT Annual Meeting Abstracts, 113(S1), pp. S5–S100. Available at: https://ascpt.onlinelibrary.wiley.com/doi/abs/10.1002/cpt.2835.

Kareva, I. & Karev, G., 2018. From Experiment to Theory: What Can We Learn from Growth Curves? Bulletin of mathematical biology, 80(1), pp. 151–174.

Kareva, I., Zutshi, A. & Kabilan, S., 2018. Guiding principles for mechanistic modeling of bispecific antibodies. Progress in Biophysics and Molecular Biology, 139, pp. 59–72.

Kreutz, C., Raue, A. & Timmer, J., 2012. Likelihood based observability analysis and confidence intervals for predictions of dynamic models. BMC Systems Biology, 6(1), pp. 1–9.

Kreutz, C. & Timmer, J., 2009. Systems biology: experimental design. The FEBS journal, 276(4), pp. 923–942.

Kutalik, Z., Cho, K.-H. & Wolkenhauer, O., 2004. Optimal sampling time selection for parameter estimation in dynamic pathway modeling. Biosystems, 75(1-3), pp. 43–55.

Lindauer, A. et al., 2017. Translational pharmacokinetic/pharmacodynamic modeling of tumor growth inhibition supports dose-range selection of the anti–PD-1 antibody pembrolizumab. CPT: pharmacometrics & systems pharmacology, 6(1), pp. 11–20.

Luo, M.C., Nikolopoulou, E. & Gevertz, J.L., 2022. From fitting the average to fitting the individual: A cautionary tale for mathematical modelers. Frontiers in Oncology, p. 1311.

Malmborg, A.-C. & Borrebaeck, C.A., 1995. BIAcore as a tool in antibody engineering. Journal of immunological methods, 183(1), pp. 7–13.

Marjoram, P., 2013. Approximation bayesian computation. OA genetics, 1(3), p. 853.

Muller, P.Y. et al., 2009. The minimum anticipated biological effect level (MABEL) for selection of first human dose in clinical trials with monoclonal antibodies. Current opinion in biotechnology, 20(6), pp. 722–729.

Muñoz-Tamayo, R. et al., 2018. To be or not to be an identifiable model. Is this a relevant question in animal science modelling? Animal, 12(4), pp. 701–712.

Nocedal, J., Öztoprak, F. & Waltz, R.A., 2014. An interior point method for nonlinear programming with infeasibility detection capabilities. Optimization Methods and Software, 29(4), pp. 837–854. Available at: https://www.mathworks.com/help/optim/ug/constrained-nonlinear-optimization-algorithms.html.

Olofsen, E., Dinges, D.F. & Van Dongen, H., 2004. Nonlinear mixed-effects modeling: individualization and prediction. Aviation, space, and environmental medicine, 75(3), pp. A134–A140.

Owen, J.S. & Fiedler-Kelly, J., 2014. Introduction to population pharmacokinetic/pharmacodynamic analysis with nonlinear mixed effects models, John Wiley & Sons.

Qian, G. & Mahdi, A., 2020. Sensitivity analysis methods in the biomedical sciences. Mathematical biosciences, 323, p. 108306.

Rajakaruna, H. & Ganusov, V.V., 2022. Mathematical Modeling to Guide Experimental Design: T Cell Clustering as a Case Study. Bulletin of Mathematical Biology, 84(10), p. 103.

Raue, A. et al., 2010. Identifiability and observability analysis for experimental design in nonlinear dynamical models. Chaos: An Interdisciplinary Journal of Nonlinear Science, 20(4), p. 045105.

Raue, A. et al., 2013. Lessons learned from quantitative dynamical modeling in systems biology. PloS one, 8(9), p. e74335.

Raue, A. et al., 2009. Structural and practical identifiability analysis of partially observed dynamical models by exploiting the profile likelihood. Bioinformatics, 25(15), pp. 1923–1929.

Saber, H. et al., 2017. An FDA oncology analysis of CD3 bispecific constructs and first-in-human dose selection. Regulatory Toxicology and Pharmacology, 90, pp. 144–152.

Sager, J.E. et al., 2015. Physiologically based pharmacokinetic (PBPK) modeling and simulation approaches: a systematic review of published models, applications, and model verification. Drug Metabolism and Disposition, 43(11), pp. 1823–1837.

Sher, A. et al., 2022. A Quantitative Systems Pharmacology perspective on the importance of parameter identifiability. Bulletin of Mathematical Biology, 84(3), pp. 1–15.

Simeoni, M. et al., 2004. Predictive pharmacokinetic-pharmacodynamic modeling of tumor growth kinetics in xenograft models after administration of anticancer agents. Cancer research, 64(3), pp. 1094–1101.

Steiert, B. et al., 2012. Experimental design for parameter estimation of gene regulatory networks. PloS one, 7(7), p. e40052.

Tiwari, A. et al., 2016. Assessing the impact of tissue target concentration data on uncertainty in in vivo target coverage predictions. CPT: pharmacometrics & systems pharmacology, 5(10), pp. 565–574.

Tiwari, A. et al., 2017. Optimal Affinity of a Monoclonal Antibody: Guiding Principles Using Mechanistic Modeling. The AAPS journal, 19(2), pp. 510–519.

Vafa, O. & Trinklein, N.D., 2020. Perspective: designing T-cell engagers with better therapeutic windows. Frontiers in oncology, 10, p. 446.

Wieland, F.-G. et al., 2021. On structural and practical identifiability. Current Opinion in Systems Biology, 25, pp. 60–69.

Wu, H. et al., 2008. Parameter identifiability and estimation of HIV/AIDS dynamic models. Bulletin of mathematical biology, 70, pp. 785–799.

Zhang, J. et al., 2022. A Phase 1b Adaptive Androgen Deprivation Therapy Trial in Metastatic Castration Sensitive Prostate Cancer. Cancers, 14(21), p. 5225.

Zhang, X.-Y. et al., 2015. Sobol sensitivity analysis: a tool to guide the development and evaluation of systems pharmacology models. CPT: pharmacometrics & systems pharmacology, 4(2), pp. 69–79.

Zhao, P. et al., 2011. Applications of physiologically based pharmacokinetic (PBPK) modeling and simulation during regulatory review. Clinical Pharmacology & Therapeutics, 89(2), pp. 259–267.

Zi, Z., 2011. Sensitivity analysis approaches applied to systems biology models. IET systems biology, 5(6), pp. 336–346.

